# Automagic: Standardized Preprocessing of Big EEG Data

**DOI:** 10.1101/460469

**Authors:** Andreas Pedroni, Amirreza Bahreini, Nicolas Langer

**Author notes:** corresponding authors: Andreas Pedroni, Nicolas Langer, Address: Psychologisches Institut, Methoden der Plastizitätsforschung, Andreasstrasse 15, AND 4.58, CH-8050 Zürich, Switzerland.

## Abstract

Electroencephalography (EEG) recordings have been rarely included in large-scale studies. This is arguably not due to a lack of information that lies in EEG recordings but mainly on account of methodological issues. In many cases, particularly in clinical, pediatric and aging populations, the EEG has a high degree of artifact contamination and the quality of EEG recordings often substantially differs between subjects. Although there exists a variety of standardized preprocessing methods to clean EEG from artifacts, currently there is no method to objectively quantify the quality of preprocessed EEG. This makes the commonly accepted procedure of excluding subjects from analyses due to exceeding contamination of artifacts highly subjective. As a consequence, P-hacking is fostered, the replicability of results is decreased, and it is difficult to pool data from different study sites. In addition, in large-scale studies, data are collected over years or even decades, requiring software that controls and manages the preprocessing of ongoing and dynamically growing studies. To address these challenges, we developed Automagic, an open-source MATLAB toolbox that acts as a wrapper to run currently available preprocessing methods and offers objective standardized quality assessment for growing studies. The software is compatible with the Brain Imaging Data Structure (BIDS) standard and hence facilitates data sharing. In the present paper we outline the functionality of Automagic and examine the effect of applying combinations of methods on a sample of resting EEG data. This examination suggests that applying a pipeline of algorithms to detect artifactual channels in combination with Multiple Artifact Rejection Algorithm (MARA), an independent component analysis (ICA)-based artifact correction method, is sufficient to reduce a large extent of artifacts.

## 1 Introduction

Major breakthroughs in translating insights from research in neuroscience to diagnose or treat mental disorders have been rare (Kapur, Phillips, and Insel 2012). To foster advances, several suggestions have been made to change paradigms and practices in research. This includes reconceptualizing the definition of mental disorders with concepts that are closer to their biological underpinnings and searching for diagnostic markers at this level (Cuthbert and Insel 2013). Furthermore, it has been suggested to move away from studying disorders in isolation and rather investigate disorders in the natural co-occurence of multiple disorders. At last, it is proposed that study samples should cover a heterogeneous composition of symptoms in various degrees of severity, facilitating the generalization and application of results to broader populations (Woo et al. 2017).

To implement these suggestions, studies with vast sample sizes are needed. This has become manifest in an increasing number of large-scale research initiatives (Eickhoff et al. 2016), such as the IMAGEN study (Schumann et al. 2010), the UK Biobank (Sudlow et al. 2015), the Autism Brain Imaging Data Exchange (Di Martino et al. 2014) or the Alzheimer’s Disease Neuroimaging Initiative (ADNI) (Mueller et al. 2005). These biomarker initiatives have assessed various sets of neuropsychological, lifestyle phenotypes and brain-imaging measures, predominantly using structural and functional magnetic resonance imaging (s/fMRI). One aspect, which is at first sight surprising, is that with the exception of one notable initiative, the Healthy Brain Network study (Alexander et al. 2017; Langer, Ho, Alexander, et al. 2017), electroencephalography (EEG) measures have not been included in any large biomarker initiative.

We believe that the current paucity of EEG measures in large-scale biomarker studies is not due to a lack of information about neuronal processes that can be gained from EEG. Complementing the rather static view of the brain that is offered by MRI with information at the speed of the actual brain processes, a wealth of studies has shown that the EEG deviates in various mental disorders from healthy controls. EEG recorded during rest is highly reliable (Näpflin, Wildi, and Sarnthein 2007; Lund et al. 1995) and is altered in numerous mental diseases, such as ADHD (Boutros, Fraenkel, and Feingold 2005), autism spectrum disorders (Wang et al. 2013), schizophrenia (Boutros et al. 2008), depression and anxiety disorders (Thibodeau, Jorgensen, and Kim 2006), and dementia and mild cognitive impairment (Jelic and Kowalski 2009). There are also abundant studies showing that task-related EEG parameters differ between healthy controls and patients suffering from various disorders (Pfefferbaum et al. 1984; Cecchi et al. 2015; Duncan et al. 2009). Furthermore, specific event-related potentials are suggested to reflect endophenotypes, for instance for anxiety disorders, depression, and substance abuse (Olvet and Hajcak 2008). Task-related EEG additionally offers insights into the processing steps that are required to perform these tasks, thus allowing insights into *why* a person performs more or less well in a cognitive task (Langer, Ho, Pedroni, et al. 2017). This reflects an aspect, which complements neuropsychological testing (often included in biomarker studies), that typically only provides information about performance in the form of a single score. Finally, EEG measures are also favorable from an economic point of view: If EEG-based biomarkers are identified, the financial burden of implementing widespread screening for such markers will be low compared to MRI-based screening.

One major challenge that has hindered the large-scale application of EEG is the difficulty of obtaining EEG recordings that are comparable in their quality between subjects. The EEG signal is influenced by technical factors and by characteristics of the recorded subject. For instance, the EEG signal is influenced by environmental factors such as temperature and air humidity. These factors interact with sources of noise, such as line noise, or other sources of electromagnetic noise (Kappenman and Luck 2010). In addition, subject-related artifacts, typically reflecting unwanted physiological signals (such as eye movements, eye blinks, muscular noise, heart signals and sweating), may differ from subject to subject and may interact in a complex manner with non-physiological artifacts. The resulting artifact signals in the EEG are typically more prominent than the signal of interest (i.e. brain activity) and hence require preprocessing in the form of artifact cleaning or artifact correction (Keil et al. 2014).

Various automated artifact preprocessing pipelines have been developed to preprocess EEG data. They apply methods of source separation techniques (Delorme, Sejnowski, and Makeig 2007; Jung et al. 2000), regression methods, and linear decomposition (Parra et al. 2005) to clean EEG data. In addition, there exist various methods of detecting and removing artifactual channels and segments from EEG (Nolan, Whelan, and Reilly 2010; Bigdely-Shamlo et al. 2015). A number of processing pipelines combine parts of these methods and/or extend them. The PREP pipeline (Nima Bigdely-Shamlo et al. 2015) offers standardized early-stage EEG processing, includes sophisticated methods to remove line-noise (using Clean Line, Mullen 2012) and robust average referencing by iteratively detecting and interpolating artifactual channels. The Harvard Automated Processing Pipeline for EEG (HAPPE) (Gabard-Durnam et al. 2018) reflects a standardized pipeline that additionally offers artifact correction using Wavelet-enhanced independent component analysis (ICA), ICA and Multiple Artifact Rejection Algorithm (MARA, Winkler, Haufe, and Tangermann 2011; Winkler et al. 2014). The Computational Testing for Automated Preprocessing (CTAP) toolbox (Cowley, Korpela, and Torniainen 2017; Cowley and Korpela 2018) has a similar goal but additionally offers functions to optimize the different preprocessing methods by allowing the user to compare different pipelines. More recently, the Batch Electroencephalography Automated Processing Platform (BEAPP, Levin et al. 2018) reflects a platform for batch processing of EEG that allows the creation of preprocessing pipelines with a variety of options that can be re-applied to new datasets, hence facilitating the exact methodological replication of studies.

Although all of these preprocessing pipelines do a remarkably good job in removing artifacts from EEG, arguably none of them would claim to be able to do this perfectly. Therefore, the extent to which a subject produces artifacts, such as moving the eyes or blinking, not only influences the signal in the raw EEG but may leave spurious traces in the preprocessed EEG. As a consequence, preprocessed EEG may vary to large extent between subjects. In some cases where the EEG signal is highly contaminated with noise, which is often the case in heterogeneous study populations such as developmental studies with children and elderly subjects or in clinical studies, artifact correction cannot recover the EEG signal from the noise in a satisfactory way. It is then often unavoidable common practice to dismiss a selection of EEG recordings due to inferior quality from the analyses that enter a study.

The decision on whether a recording has acceptable quality to be submitted for further analysis is highly subjective and depends on many factors such as the recording length (i.e. number of trials), sample size, the effect sizes of interest and, foremost, the specific research question. In studies with small sample sizes it is manageable for one researcher to perform this classification, which is typically based on the visual inspection of the preprocessed EEG^1^. Even in the case that one person assesses an entire dataset, the classification within the dataset might vary as a function of his/her motivation, tiredness and depend on the previously reviewed datasets. Moreover, it can be highly problematic if the data are not blinded and are classified with knowledge about the hypotheses. Classification may become more intricate in large-scale studies with thousands of datasets, where it is not feasible for one person to classify all data sets. The classification sensitivity may also change from researcher to researcher, and thus may introduce biases in the study sample that lower the generalizability of results.

The recently introduced pipelines HAPPE (Gabard-Durnam et al. 2018) and CTAP (Cowley, Korpela, and Torniainen 2017; Cowley and Korpela 2018) offer standardized metrics of the data quality allowing objective classification of datasets. For HAPPE, quality metrics are the number of interpolated channels, the number of epochs that survive epoch-wise artifact rejection after preprocessing, the number of independent components (IC) that are rejected as artifact components, and the retained variance of the EEG data after IC rejection. CTAP offers the percentage of bad channels, bad segments, bad epochs and bad components. Although these metrics are helpful to assess the quality of data, they rely on how well the preprocessing was able to identify artifacts. For instance, the percentage of interpolated channels (i.e. channels that are contaminated by artifacts) depends on how well a bad-channel detection algorithm is capable of detecting noisy channels. Since the noise may vary between subjects, the bad-channel detection algorithm may vary in the sensitivity to identify a noisy channel. In addition, the retained variance of the EEG data after IC rejection is influenced by how clean the data are when entering the ICA. More retained variance does not per se indicate less noisy data. Therefore, the quality metrics are only valid if the preprocessing is equally capable of identifying artifacts in each recording. This is an assumption on which our experience casts doubt.

A range of alternative quality metrics (that are not included in toolboxes), are based on the ratio of signal and noise in specific paradigms. Some quality metrics have been proposed that are related to the amplitude or spectral power of certain ERP events or time-windows (Oliveira et al. 2016; Zander et al. 2017; Chi et al. 2012; Ries et al. 2014; Fiedler et al. 2015). For resting state EEG, signal-to-noise has been quantified by means of the extent of the alpha block, or frontal theta in eyes-closed versus eyes-open conditions (Radüntz 2018) or more generally the alpha band compared to the PSD ratio (Liu et al. 2019). While these measures are of great use to quantify the quality of the signal of interest in a dataset, they are very specific to a research question and it is difficult to compare the quality of datasets with different paradigms. Furthermore, the quality of the signal not only depends on the quality of the data but also on the number of trials, respectively the duration of the recording.

We propose alternative quality metrics that are based on the quantification of the absolute signal strength, a method akin to the common practice of selecting trials/epochs of good quality by applying absolute µ*V* thresholds (Kaibara, Holloway, and Young 2010; Nakamura et al. 2005). This set of metrics is generally applicable and does not dependent on the identification of artifacts or bad channels and hence offers more standardized inclusion/exclusion criteria for EEG recordings.

A further practical problem that hinders the implementation of EEG measurements in large-scale studies is a lack of software that allows the handling of large and, in particular, growing datasets. Large-scale studies and longitudinal studies typically run over many years and it is impractical to hold back data preprocessing until the end of a study. Often, research questions can be addressed in an *a priori* defined subsamples of a large study and therefore scientific progress would be slowed down if data are not released until the endpoint of a study. This requires that preprocessing software is capable of handling growing studies by keeping track of the already preprocessed datasets and by freezing preprocessing settings (and software versions) to assure a standardized procedure.

In addition to the urge of standardizing preprocessing, there is a need for a principled format to organize, harmonize, and share data. In the recent years, EEG datasets have been made increasingly openly available (for a list of shared EEG datasets, see https://github.com/voytekresearch/OpenData) and it has been shown that integrating EEG datasets across studies offers unique insights unavailable in single studies (Bigdely-Shamlo, Touryan, et al. 2018; Bigdely-Shamlo, Touyran, et al. 2018). The principled way of data sharing has been successfully adopted in the domain of MRI data with the introduction of the Brain Imaging Data Structure (BIDS), the emerging standard for the organisation of neuroimaging data (Gorgolewski et al. 2016). Various extensions of the BIDS format (including extensions for EEG data (Pernet, Appelhoff, et al. 2018)) have been proposed (https://bids.neuroimaging.io) that not only provide a standard for the respective data modality but moreover facilitate the integration between data of different modalities (e.g. simultaneous fMRI and EEG recordings). To make EEG data sharing simple and intuitive, it would therefore be beneficial if a preprocessing pipeline supports BIDS format as in- and output.

In the present paper we aim to narrow the current gaps in EEG preprocessing that hamper large-scale studies and open science practices, by presenting a software solution that implements a selection of currently available artifact correction procedures in addition to a novel quality assessment tool that relies on standardized and quantitative quality metrics that are independent from the preprocessing. Beyond this, the software handles growing studies, supports the BIDS format and hence facilitates data sharing. To foster scientific transparency and make exact methodological replications of studies easier, detailed information of all processing steps are stored in a log-file in accordance to (Pernet, Garrido, et al. 2018) and project settings of already existing studies can be saved and imported. The software, called Automagic, is based on Matlab, open source and freely available under the terms of the GNU General Public License. It is available from https://github.com/methlabUZH/automagic. In the remainder of this manuscript, we first outline Automagic’s functionalities, then describe how Automagic is embedded in the current landscape of preprocessing software and provide a validation of applying combinations of methods on a sample of resting EEG data.

## 2 The Automagic processing pipeline for big EEG data

### 2.1 Overview

Automagic has been developed with the intention of offering a user-friendly preprocessing software for big (and small) EEG datasets. One major focus of the software lies in the standardization of the preprocessing and quality control and the transparent and standardized reporting to facilitate open science practices. The software can be fully run by command line or with a graphical user interface (GUI) and therefore does not require any knowledge of programming. It runs in MATLAB (from Version 2016b) and can be installed easily. Due to the fact that Automagic is built on several MATLAB functions that are compatible with EEGLAB (Delorme and Makeig 2004), one of the most used open-source frameworks for EEG analysis), more experienced users can readily modify Automagic to their specific needs. In addition, Automagic can be used as an EEGLAB extension.

Automagic contains three basic sections, the *project section*, the *configuration section*, and the *quality assessment section*. A typical workflow involves the following steps: In the project section, the user defines the input and output folders of the processing and acquisition information about the EEG data, such as the format, channel locations, and electrooculogram (EOG)-channels. The configuration section allows the selection of various preprocessing steps that will be applied to the EEG. After preprocessing of the data and the interpolation of bad channels, the user can review the quality of the preprocessing in the quality section. Here, EEG datasets can be displayed as heatmaps (with dimensions of channels and time) or as classical EEG channel plots (using the eegplot function of EEGLAB), allowing the user to obtain a comprehensive overview of the data quality. In addition, various quality criteria are computed such as the ratio time windows of high voltage amplitudes. Finally, the user can define cutoffs for these quality criteria to objectively include datasets of good quality and exclude datasets of too poor quality. For each dataset, the exact sequence of the preprocessing steps is stored in a log-file and all parameters of the preprocessing and the quality measures are stored in a MATLAB structure and in a JSON-file that is stored along with the preprocessed data. The entire project can be exported to BIDS format, which includes a raw data folder (with the raw data), a derivative folder with the preprocessed data and the code that translates raw to derivative. In the following sections, we will walk through the processing steps (see Figure 1 for an overview) of Automagic and provide detailed information.

**Figure 1.**
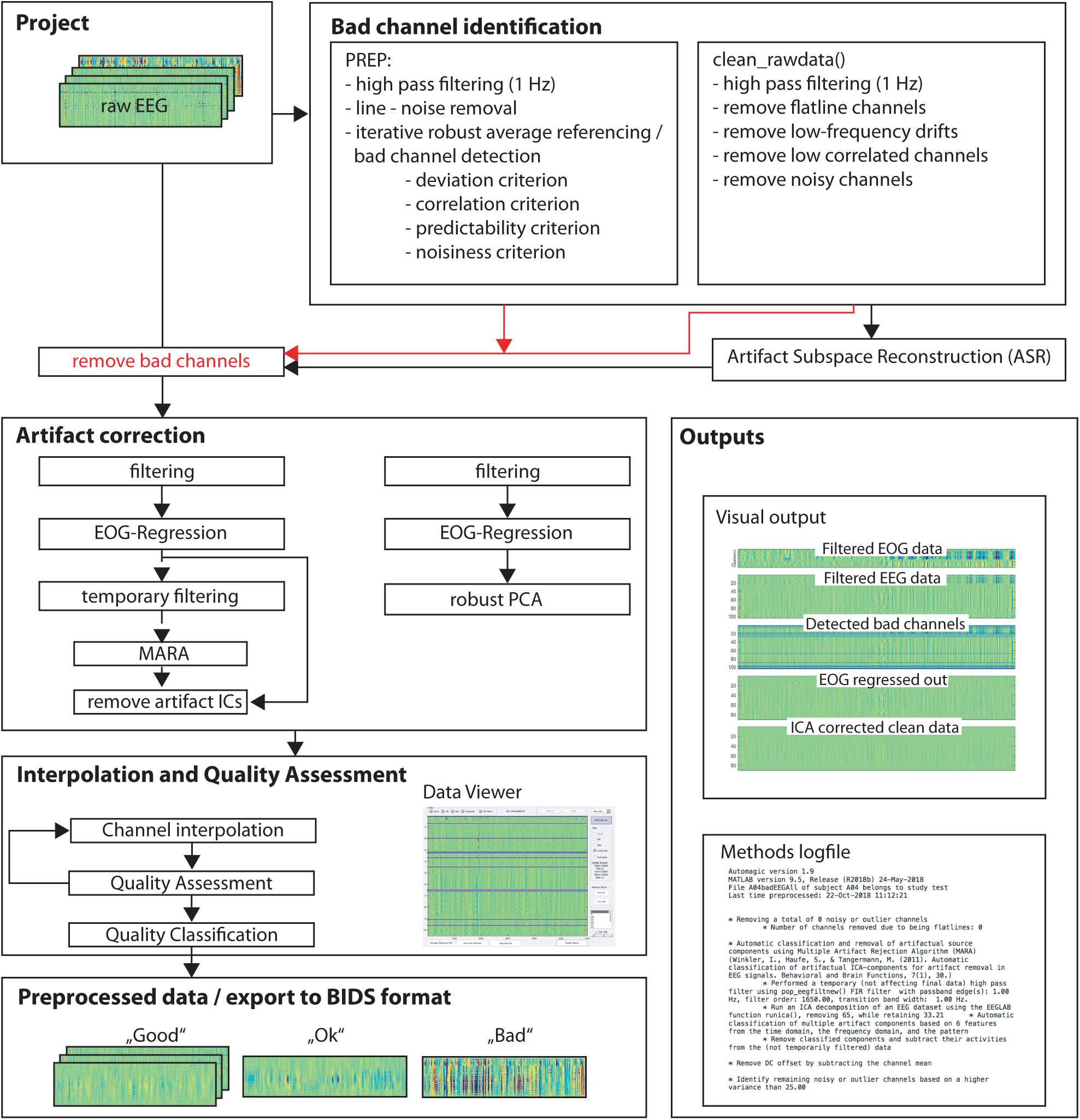
Typical workflow of Automagic. The raw EEG data are organized in a project. Each dataset within a project enters the bad channel identification routines chosen by the user, detecting noisy, flat or outlier channels. These channels (red arrow) are subsequently removed from the original EEG. When choosing Artifact Subspace Reconstruction (ASR), the EEG is additionally corrected for corrupt segments at this point. In a next step, the EEG can be filtered and the user can choose to apply the artifact correction methods of EOG-regression and MARA or rPCA. In a next step, the bad channels are interpolated and the quality of each dataset is assessed using a number of objective criteria. Finally, the user can categorize the quality of each dataset into “Good”, “OK” and “Bad”, by applying cutoffs to the quality criteria. In addition, a visual summary of the preprocessing for each dataset and a log-file with the applied methods are saved.

#### 2.1.1 Project

The basic idea behind a *project* is that the user defines a *data folder,* which is the source of preprocessing, along with a *results folder,* which is the target for the preprocessed data, additional output files and contains a list of configurations for the preprocessing that is stored in the results folder in a so-called *project file (project_state.mat)*. Once the configurations have been defined and a project has been created, all the configurations are fixed and can no longer be changed, thus assuring that all datasets are processed in an identical fashion. For each project the versions of Automagic and Matlab are saved in the log files and also in the actual output files. On https://github.com/methlabUZH/automagic all available versions of Automagic will be accessible. In addition, researchers can build a software container (e.g. Docker), stored in a repository. This will guarantee that preprocessing can be exactly replicated. A tutorial how to build a Docker container can be found on the GitHub Wiki page: https://github.com/methlabUZH/automagic/wiki/Create-Docker-Image. In contrast to having the configurations fixed, the input files are not; hence, it is always possible to add (or to delete) datasets to a data folder and Automagic detects these changes whenever the application is restarted. This guarantees that all EEG files are preprocessed in a standardized way.

#### 2.1.2 EEG data import

Automagic supports most common EEG file types, specifically all the EEG file types that can be imported with EEGLAB (see https://sccn.ucsd.edu/wiki/EEGLAB_Extensions_and_plug-ins). In addition, the EGI HCGSN “raw“ file format is supported for 128 and 256 electrode data sets (or 129 and 257 if reference electrode is included respectively), with included channel locations.

### 2.2 Bad channel identification

In a first step, so-called “bad channels” are identified^2^. Generally, bad channels are those with low recording signal-to-noise ratio (SNR) or very low or no signal throughout a considerable time span of the EEG recording. If there are only few channels affected by such noise/low signal, it is impractical to reject/correct entire time windows of the EEG for this reason. Instead, in many cases it is more sensible to interpolate the signal of bad channels by the combined signal of channels with adequate SNR or to discard bad channels for further analyses.

Automagic creates a copy of the original raw EEG data, which is submitted to algorithms whose purpose is to detect bad channels. In this way, the original data are unaffected by the high-pass filtering with a cutoff of 1 Hz that is performed by default for the currently supported bad channel removal algorithms. Note that the bad channel identification procedures only *detect* bad channels and do *not* modify (e.g. interpolate channels) the EEG data. Bad channels are interpolated after the preprocessing.

#### 2.2.1 Bad channel identification and robust average referencing using the PREP pipeline

By default, Automagic uses the PREP pipeline (Bigdely-Shamlo et al. 2015) that offers four sophisticated bad channel identification algorithms and a method to estimate a robust average reference. The specific processing steps are described in detail in (Bigdely-Shamlo et al. 2015) and on http://vislab.github.io/EEG-Clean-Tools/. Here, we only outline the key steps. In a first step, the copy of the original EEG data is high-pass filtered with a cutoff of 1 Hz. Next, power line noise is removed, using the “cleanLine” EEGLAB plugin developed by Mullen (2012), which estimates the amplitude and size of a deterministic sinusoid at a specified frequency embedded in locally white noise. The algorithm requires approximate indications of the frequencies to be removed, which are by default multiples from the frequency of the line power filter input, up to half of the Nyquist frequency.

After these two initial steps, PREP iteratively estimates a robust average reference. The robust average reference method subtracts the average of all “good” (i.e. not “bad”) channels from each channel. Specifically, the algorithm iteratively detects and interpolates bad channels to arrive at an average reference that is not affected by artifacts. It uses four measures for this: extreme amplitudes (deviation criterion), lack of correlation with any other channel (correlation criterion), lack of predictability by other channels (predictability criterion), and unusual high frequency noise (noisiness criterion). Note that the estimated robust average reference is only computed but not applied to the EEG data. Therefore, the data remain referenced to the initial reference. Note that the PREP pipeline is fairly computationally expensive, especially on datasets with many EEG channels, and it is advisable to allow PREP to use the parallel processing offered by MATLAB, which is done by default.

#### 2.2.2 Bad channel identification using clean_rawdata()

Computationally less expensive and faster alternatives (or an addition) to the bad channel identification algorithms offered by PREP, are the three bad channel identification algorithms of the EEGLAB plugin clean_rawdata() (http://sccn.ucsd.edu/wiki/Plugin_list_process). The first algorithm detects channels that have no signal variation for a duration of longer than 5 s. The EEG of the remaining channels is then filtered with a forward–backward (non-causal) filter (with transition band of 0.5 - 1 Hz and stop-band attenuation of 80 db) to remove slow-wave drifts. The second algorithm (equivalent to the predictability criterion of PREP) searches for channels with a lower correlation to its robust estimate (based on other channels) than a given threshold. Specifically, (using the default settings) it calculates the correlation of each channel in a time window of 5 s of the EEG to its random sample consensus (RANSAC) reconstruction (with 50 samples derived from a subset of 25% of all channels). If the correlation is below 0.85 in more than 40% of times, the respective channel is identified as bad. The third algorithm removes EEG channels with excessive line noise. It extracts the signal above 50 Hz from the raw EEG, which is considered line noise. Then the median absolute deviation (MAD) of the difference between raw EEG and the line noise is calculated and divided by the MAD of the line noise, which gives a relative noise-to-signal estimate for each channel. This estimate is z-transformed and channels with a z-value above 4 (i.e. 4 standard deviations above the channel population mean) are marked as bad channels.^3^

In addition to the bad channel identification algorithms, clean_rawdata() offers a function that automatically detects and repairs artifacts using the Artifact Subspace Reconstruction (ASR) method (Mullen et al. 2013). This method does not identify bad channels, but detects “bad” time windows (with pronounced bursts of noise) in the data and attempts to interpolate this data based on the rest of the EEG signal during these time periods. A second function identifies and removes time periods that could not be recovered by ASR (with abnormally high power from the data). Note that the selection of these two additional processing steps has consequences for the original EEG as the EEG dataset might be shortened by the time periods that could not be adequately recovered. In a limited set of tests with small data sets, we observed that the time periods rejected by ASR varied and hence were not perfectly replicable. Therefore, we suggest testing the use of ASR with respective data sets.

#### 2.2.3 Residual bad channel detection

In addition to these algorithms, we found it for some datasets helpful to identify channels that show exceedingly high variance over time. Bad channels are identified by searching for channels with a standard deviation exceeding a certain threshold (by default 25 µV). Note that this residual channel identification algorithm is applied after the entire preprocessing. The reason for applying this step after preprocessing is to identify whether residual noisy channels also remain after the artifact correction, which we have observed in some cases.

#### 2.2.4 Bad channel removal

After having run either (or both) of the bad channel identification pipelines (clean_rawdata() and/or the PREP pipeline) the selection of identified bad channels is excluded from the original EEG data (see Fig. 1). This is done because, for instance, the ICA used in a later processing step in many cases does not work with bad channels (e.g. flat channels). Note that from this point on, the downstream processing changes the EEG signals but up to this point channels are interpolated.

### 2.3 Filtering

In this step, EEG data can be filtered using low-pass, high-pass and or notch-filter, implemented in EEGLAB’s pop_eegfiltnew() function.

### 2.4 Artifact correction

Although bad channel identification methods have the goal of identifying a distinct set of channels with low SNR throughout a considerable time span of the EEG recording, many artifacts span multiple channels, while being more temporally isolated, such as eye-blinks and movement muscle artifacts. One simple approach to handle such artifacts is to detect them (e.g. EOG data with amplitudes of ±75 µV) and delete these time segments. However, this can potentially lead to a large loss of data, consequently reducing the SNR and the quality of downstream analyses (i.e. too few trials remain for an ERP analysis). To avoid data loss, various methods exist that remove the artifacts while preserving the underlying neural activity. In Automagic three such artifact correction methods are implemented using the EOG regression, MARA, an ICA-based correction method, and a robust principal component analysis (rPCA) method.

#### 2.4.1 EOG regression (EOGr)

In this step, artifacts that are caused by eye movements are separated from the EEG signal using the subtraction method that relies on linear transformation of the EEG activity (see Parra et al. 2005). This method is similar to the classical regression approach, where EOG electrodes are used as a reference signal that is subtracted in proportion to their contribution to each EEG channel (Croft and Barry 2000). The main difference to the classical approach is that EEG channels are also used to construct an optimal estimate of the EOG activity as reference for subtraction.

#### 2.4.2 Multiple Artifact Rejection Algorithm (MARA)

The Multiple Artifact Rejection Algorithm (MARA) (Winkler, Haufe, and Tangermann 2011; Winkler et al. 2014), used as a default, is an open-source EEGLAB plug-in, which automatizes the process of hand-labeling independent components (ICs) for artifact rejection. In short, ICs of the EEG are separated by independent component analysis (ICA) using the runica() function implemented in EEGLAB. MARA classifies ICs into those that reflect artifacts and those that reflect EEG. This classification is based on a supervised learning algorithm that has learned from expert ratings of 1290 components considering six features from the spatial, the spectral and the temporal domain. The ICs that are classified as artifacts are removed and their activities are subtracted from the data. Note that prior to the ICA, a copy of the data is made and high-pass filtered with a cutoff of 1 Hz. This is done because the ICs classified as artifacts are removed from the original data, allowing the data to be filtered with lower cutoff frequencies than those optimal for running the ICA in MARA.

#### 2.4.3 Noise removal using robust PCA (rPCA)

An alternative (and faster) way of removing artifacts is the use of robust principal component analysis (rPCA). The algorithm (Lin, Chen, and Ma 2010) estimates a low-dimensional subspace of the observed data, reflecting the signal that is cleaned from sparse corruptions. In contrast to the classical PCA, the rPCA is able to efficiently estimate the true low-dimensional subspace even when the observations are grossly corrupted. We advise using the ICA instead of the rPCA, as previously suggested by Jung and colleagues (2000) because we have observed that in some cases oscillatory signals that are for example induced in steady state visual evoked potential (SSVEP) paradigms are removed by the rPCA.

### 2.5 DC offset removal

As a final preprocessing step, the DC offset can be optionally removed by subtracting the channel-wise mean from each data point. Although DC offset removal is in most situations unnecessary, when a high-pass filter is applied, we offer the researcher the option of applying a DC offset removal. Several papers have addressed the issue of inappropriate usage of high-pass filters, which can produce artifactual effects. The authors provide guidelines on how to minimize filtering artifacts (e.g. Tanner et al., 2015, Acunzo et al., 2012). One of these guidelines recommends to compare the high-pass filtered data with only DC-offset corrected data (Acunzo et al., 2012) to evaluate the high-pass filter effects. Thus, Automagic enables the option of DC-offset removal.

### 2.6 Preprocessed data and auxiliary outputs

The preprocessed datasets are saved sequentially.^4^ Therefore, the user can interrupt and continue the preprocessing at any time without losing previously preprocessed data. Automagic saves the preprocessed data as a MATLAB MAT-file (Version 7.3). After preprocessing, the prefix *“p”* (for preprocessed) is added to the filename. The MAT-file contains a structure named *EEG* that corresponds to the EEG structure of EEGLAB (see EEGLAB https://sccn.ucsd.edu/wiki/A05:_Data_Structures). Saving the preprocessed data in an EEGLAB *EEG* structure makes it possible to readily use EEGLAB for further analyses by simply loading the MAT-files into the MATLAB workspace.^5^ In addition the MAT-file includes a structure named *automagic*, which contains all parameters of the preprocessing and quality measures. The two structures *EEG* and *Automagic* can be selectively loaded (using *load(filename,structure)*), which saves time when loading data, for instance if one wants to inspect the information in the (small in byte-size) *Automagic* structure.

For each preprocessed dataset, a jpg. file is saved (in the same folder), providing a visual representation of the effects of the applied preprocessing methods.^6^ In addition, a log-file is saved for each dataset, describing the exact order and the used parameters of the preprocessing steps. This logfile can be used to precisely communicate the preprocessing of an EEG project in a publication in accordance with the recommendation for reporting standards of the Committee on Best Practice in Data Analysis and Sharing (COBIDAS) (Pernet, Garrido, et al. 2018).

### 2.7 Data Viewer

The user can now review the data in the *Data Viewer,* where each dataset is plotted as a heatmap and bad channels are marked with blue lines allowing the user to obtain a comprehensive overview of the data quality. The preprocessed datasets can be visually compared to the raw 1 Hz high-pass filtered data. Furthermore, it is possible to manually select bad channels that have not been detected by the applied methods; however, for the sake of replicability we discourage users from doing this.

### 2.8 Bad channel interpolation

In a next step all the channels that have been detected as bad channels can be interpolated using the eeg_interp() function of EEGLAB with the “spherical” interpolation method as a default. The interpolation of bad channels is optional. If no interpolation is performed, the values in the rejected channels are replaced with NaN’s and are discarded from further analyses. After interpolation the EEG data are saved and the prefix “i” (for interpolated) is added to the filename.

### 2.9 Quality assessment

The goal of the quality assessment module is to provide objective measures of the data quality after preprocessing. Because datasets in a project may be differently contaminated by artifacts, more or less channels may have been detected and interpolated. Similarly, the artifact correction methods may have been more or less effective in cleaning the data from artifacts. This can lead to variability in data quality within a project and often requires the researcher to exclude overly contaminated datasets from downstream analyses. Obviously, the decision about which datasets to exclude from analysis leaves many degrees of freedom and hence may lead to an increase in Type I errors (Simmons, Nelson, and Simonsohn 2011) in the statistical inference of downstream analyses. Automagic offers a number of objective quality measures with which datasets can be compared within a project and objective exclusion criteria can be determined. These criteria can be used to facilitate the exact replication of a project or can be included in the pre-registration of an entire methodological workflow in an EEG analysis.

#### 2.9.1 Quality measures

*Ratio of Bad Channels (RBC).* One indicator for good quality of the data is the ratio of identified bad and consequently interpolated channels (RBC). The more channels that are interpolated the more of the signal of interest is lost, hence the worse the data quality. Assessing the extent of signal loss is not straightforward because the signal loss depends on the spatial configuration of interpolated channels. If many bad channels are located in clusters the signal loss is greater than if bad channels are spatially scattered across the scalp. Therefore, this quality measure should be considered together with the scalp distribution of interpolated channels and clustered bad channels should be considered as more negatively affecting the quality of the data than scattered bad channels.

The quality of each dataset is furthermore assessed using three types of quality measures that are based on the magnitude of the voltage amplitudes. A range of thresholds can be defined for these measures, resulting in quality measures of different sensitivity.

*Ratio of data with overall high amplitude (OHA).* The quality measure of the overall high amplitude (OHA) is defined by calculating the ratio of data points (i.e. channels *c* x timepoints *t*) that have a higher absolute voltage magnitude *v* of *x* µV,

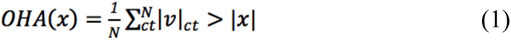

where *x* reflects a vector of voltage magnitude thresholds (e.g. *x* = 10 µV, 20 µV, 30 µV, 50 µV, 60 µV, 70 µV) and *N* reflects the number of data points. Thus, each OHA(*x*) threshold results in a quality measure that differs in its sensitivity.

*Ratio of timepoints of high variance (THV).* Similarly, the ratio of time points *t* is identified, in which the standard deviation σ of the voltage measures *v* across all channels *c* exceeds *x* µV, where *x* reflects a vector of standard deviation thresholds and *T* the number of time points. The ratio of timepoints that exceeds such a threshold reflects the timepoints of high variance (THV) criterion for a given threshold *x*:

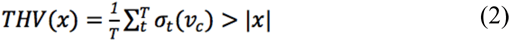

*Ratio of channels of high variance (CHV).* The same logic applies to the ratio of channels, for which the standard deviation σ of the voltage *v* measures across all time points *t* exceeds *x* µV. This ratio is reflected in the channels of high variance (CHV) criterion:

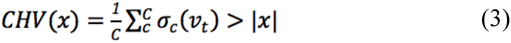

Therefore, after preprocessing and interpolating the datasets of a project, quality measures with varying sensitivity (i.e. by applying variable thresholds) for high-amplitude noise are calculated. The effect of the different thresholds is illustrated in Figure 2. The choices of the quality measure thresholds should serve to optimize the sensitivity of high amplitude noise. We found the amplitude threshold of 30 µV and the standard deviation thresholds of 15 µV to be a good starting point to quantify remaining artifacts when data are high-pass filtered with a bandpass of 0.5 Hz^7^. For data filtered with a higher bandpass, higher thresholds may be more suitable and can be changed by selecting the respective threshold in the corresponding drop-down menus in the *Quality Classification* panel.

**Figure 2.**
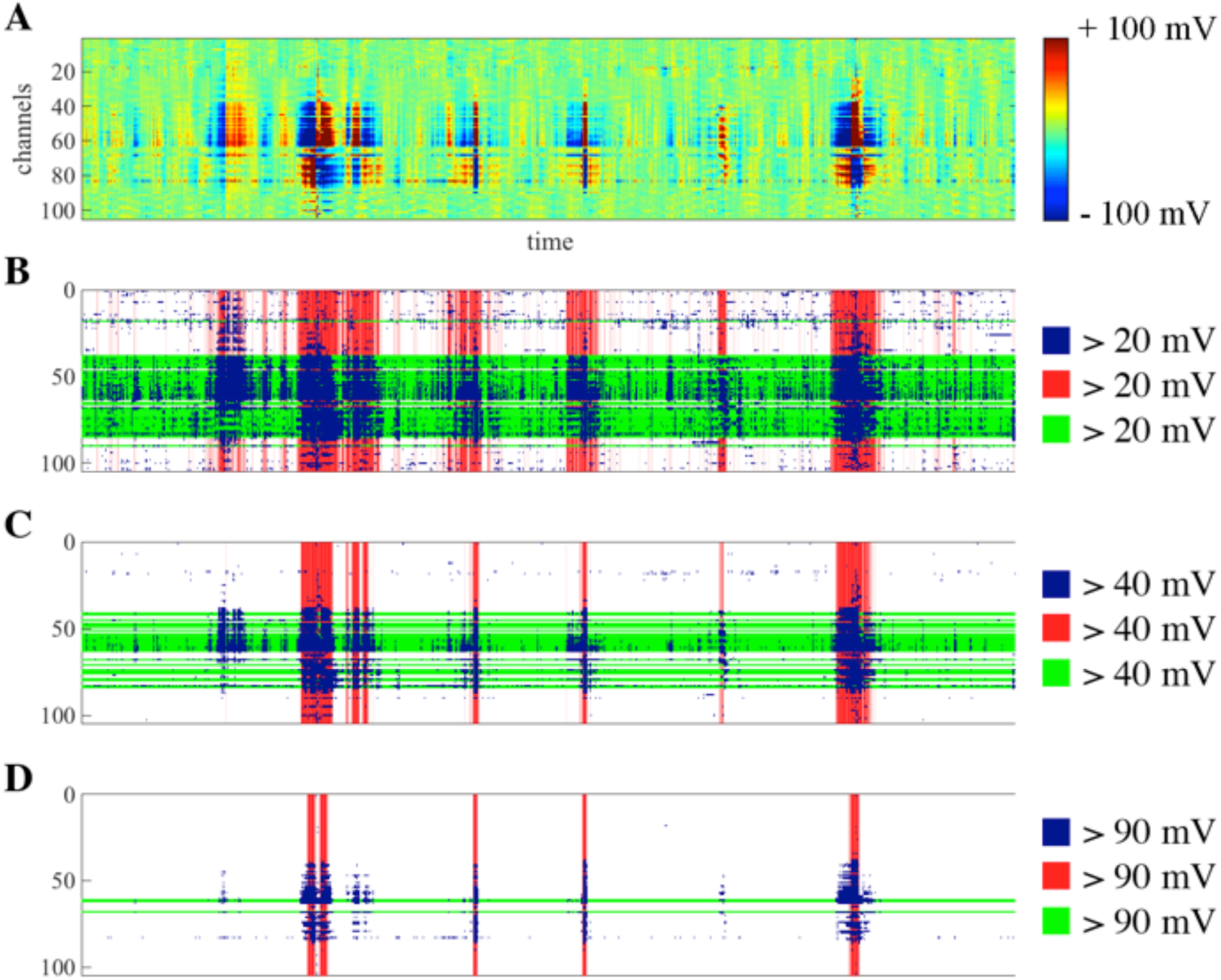
Effect of applying different thresholds to the quality measures. Panel A shows an exemplary EEG dataset as a heatmap scaled from -100 µV to +100 µV, with considerable artifacts. A) Applying rather strict thresholds of 20 µV result in large values for overall high amplitude criterion (OHA) (20%), the timepoints of high variance criterion (THV) (25%), and the channels of high variance criterion (CHV) (45%). B) and C) show the effects of applying more liberal thresholds of 40 µV and 90 µV, resulting in smaller values for OHA (8%, 2%), THV (10%, 4%), CHV (26%, 3%). Blue areas indicate data points marked by the OHA criterion, red time windows indicate epochs marked in accordance with the THV criterion, green lines indicate channels marked in accordance with the CHV criterion

We would like to emphasize that the rationale behind assessing quality with multiple measures is that the quality of a dataset strongly depends on the goal and specifics of the analysis. For instance, if a researcher is only interested in ERPs of very specific channels, he or she might ignore the CHV and RBC criterion, and only consider that the channels of interest are free of artifacts. Alternatively, if there are many trials available in a dataset, the THV criterion could be relaxed. Therefore, the user can select the thresholds and the criteria, upon which the quality assessment is based on. However, importantly, the thresholds are applied to all datasets of one project and therefore the quality standards are the same for all datasets.

#### 2.9.2 Quality classification

Using a selection of the quality measures (OHA, THV, CHV, RBC) and specified thresholds, the user can classify the datasets of one project into three categories “Good”, “OK”, or“Bad”, by applying cutoffs for each quality measure. We provide three categories for testing whether effects of interest that can be delineated in very clean EEG data (i.e. “Good” category) also hold in noisier (“OK” category) data. By equating the thresholds of “OK” to the thresholds of the “Bad” category, the data can be included/excluded in a deterministic way for later analyses.

All the default values for thresholds and category cutoffs can be modified and therefore the user can compile samples in accordance with his/her own purposes. However, to restrict oneself in repeatedly compiling categories and running analyses until effects become true (i.e. P-hacking), the user can commit to the selected categorical assignment by pressing “commit classification”, which appends the letters “g” for “Good”, “o” for “OK”, and “b” for “Bad” to the file name of the preprocessed EEG dataset. The application of such quantitative quality measures and cutoffs makes it possible to compare the ratio of “Good”, “OK”, and “Bad” categories between projects, providing insights into the comparability of the data quality of different projects/recording sites or recording apparatus and facilitate data selection for further analysis.

### 2.10 BIDS integration

Automagic can preprocess datasets in the BIDS EEG format. In addition, Automagic offers the export of a project into a brain imaging data structure (BIDS) compatible structure to facilitate data sharing. BIDS is a data sharing standard that has been originally developed for MRI data but now also encompasses other modalities, including an extension for EEG (Pernet, Appelhoff, et al. 2018). The BIDS reflects a systematic way to organize data in a folder structure with defined naming convention. An alternative standard to EEG BIDS following the same goal of enabling data sharing is the EEG Study Schema (ESS) (Bigdely-Shamlo, Makeig, and Robbins 2016). We have decided to implement BIDS EEG (instead of the ESS) because the BIDS EEG can be readily combined to other BIDS extensions such as an extension for eye tracking, or fMRI that all follow the same basic structure.

Note, that Automagic uses its own folder structure to allow an efficient management of growing datasets. Yet at any time, the user can export a project to a BIDS compatible format with a raw data folder containing the BIDS compatible data files and structures, a derivative folder with the preprocessed data and the code that performs the computations to transform data from raw to derivative. We encourage to share data using the BIDS as it facilitates communication, increases reproducibility and makes it easier to develop data analysis pipelines.

## 3 Overview of existing EEG preprocessing pipelines

A variety of more or less automated EEG preprocessing pipelines have been developed in the past years that offer methods to clean EEG data. However, after reviewing the existing pipelines we did not find a unitary solution that offered the functionality to handle BIDS format, manage growing large studies, and provide quality metrics. To outline where Automagic is positioned in the landscape of preprocessing pipelines, we have summarized functionalities and characteristics of these preprocessing pipelines in Table 1. Note that the table has been sent to the authors of the toolboxes listed in the table, so it is in agreement with them.

**Table 1.**
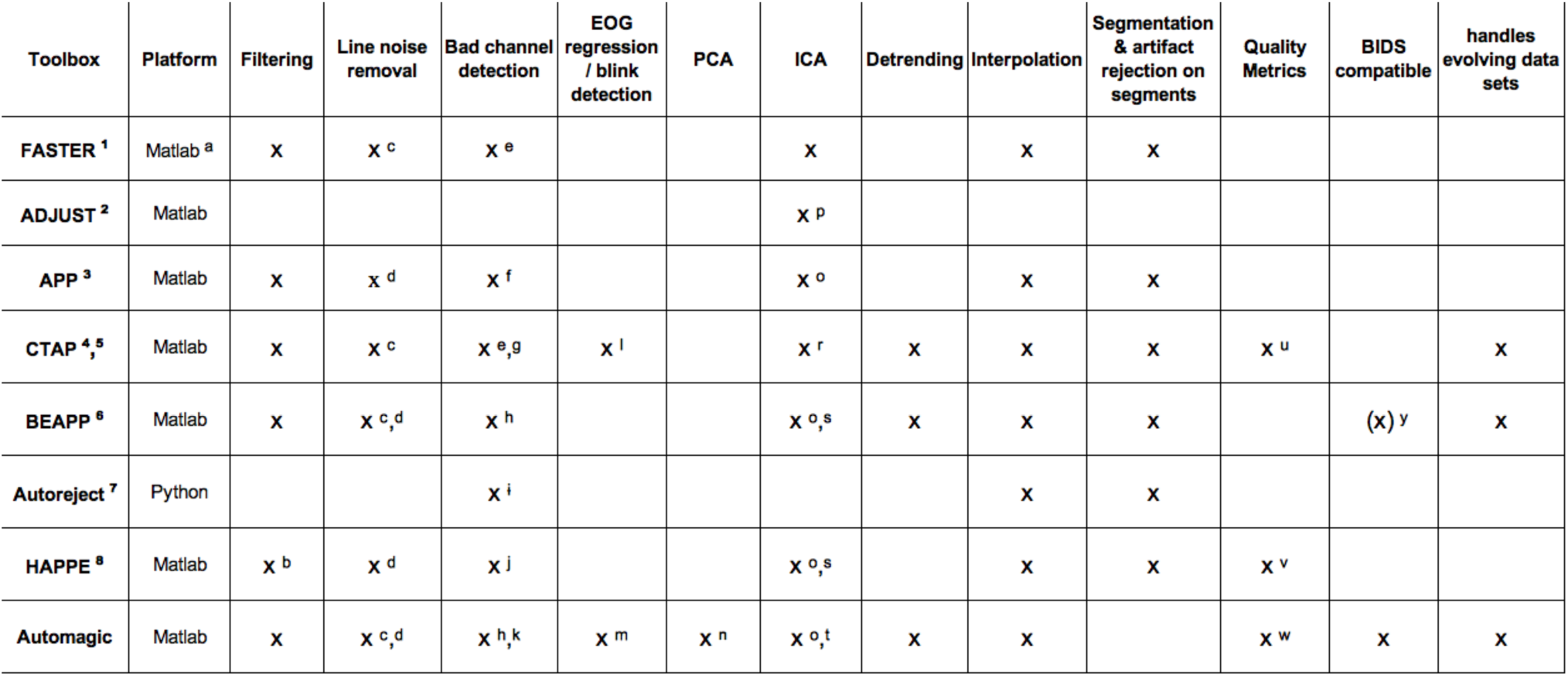
Overview of functionalities of current EEG preprocessing toolboxes. a) Python version available; b) compulsory 1 Hz high-pass; c) notch filter; d) CleanLine multi-taper regression approach; e) criteria based on variance, mean correlation & Hurst exponent; f) correlation criterion, dispersion criterion; g) criteria based on variance & channel spectra method; h) PREP: extreme amplitudes (deviation criterion), lack of correlation with any other channel (correlation criterion), lack of predictability by other channels (predictability criterion), unusual high frequency noise (noisiness criterion); i) trial wise bad sensor detection, repair or rejection; j) normed joint probability of the average log power from 1 to 125 Hz; k) clean_rawdata(): no signal variation for a duration of longer than 5 s., predictability by other channels (predictability criterion), excessive line noise; l) EOGERT blink detection; m) EOG regression; n) robust PCA; o) ICA with MARA; p) combination of stereotyped artifact-specific spatial and temporal features; r) amplitude threshold metrics from FASTER & ADJUST; s) Wavelet-enhanced ICA; t) IC Label in developmental version; u) % of bad channels, % of bad segments, % of bad epochs, % of bad components; v) number of interpolated channels, the number of epochs that survive epoch-wise artifact rejection after preprocessing, the number of independent components that are rejected as artifact components, the retained variance; w) ratio of bad channels (RBC), ratio of data with overall high amplitude (OHA), ratio of timepoints of high variance (THV), ratio of channels of high variance (CHV); y) planned to be implemented in the future. 1) Nolan, et al., (2010); 2) Mognon, et al., (2011); 3) da Cruz, et al. (2018); 4) Cowley et al. (2018); 5) Cowley, et al. (2017); 6) Levin et al. (2018); 7) Jas et al. (2017); 8) Gabard-Durnam et al. (2018).

| Toolbox | Platform | Filtering | Line noise removal | Bad channel detection | EOG regression / blink detection | PCA | ICA | Detrending | Interpolation | Segmentation & artifact rejection on segments | Quality Metrics | BIDS compatible | handles evolving data sets |
| --- | --- | --- | --- | --- | --- | --- | --- | --- | --- | --- | --- | --- | --- |
| FASTER <sup>1</sup> | Matlab <sup>a</sup> | x | x <sup>c</sup> | x <sup>e</sup> |  |  | x |  | x | x |  |  |  |
| ADJUST <sup>2</sup> | Matlab |  |  |  |  |  | x <sup>p</sup> |  |  |  |  |  |  |
| APP <sup>3</sup> | Matlab | x | x <sup>d</sup> | x <sup>f</sup> |  |  | x <sup>o</sup> |  | x | x |  |  |  |
| CTAP <sup>4, 5</sup> | Matlab | x | x <sup>c</sup> | x <sup>e, g</sup> | x <sup>l</sup> |  | x <sup>r</sup> | x | x | x | x <sup>u</sup> |  | x |
| BEAPP <sup>6</sup> | Matlab | x | x <sup>c, d</sup> | x <sup>h</sup> |  |  | x <sup>o, s</sup> | x | x | x |  | (x) <sup>y</sup> | x |
| Autoreject <sup>7</sup> | Python |  |  | x <sup>i</sup> |  |  |  |  | x | x |  |  |  |
| HAPPE <sup>8</sup> | Matlab | x <sup>b</sup> | x <sup>d</sup> | x <sup>j</sup> |  |  | x <sup>o, s</sup> |  | x | x | x <sup>v</sup> |  |  |
| Automagic | Matlab | x | x <sup>c, d</sup> | x <sup>h, k</sup> | x <sup>m</sup> | x <sup>n</sup> | x <sup>o, t</sup> | x | x |  | x <sup>w</sup> | x | x |

Most preprocessing pipelines enable bad channel detection (and interpolation), filtering (note that the Harvard Automated Processing Pipeline for EEG (HAPPE) (Gabard-Durnam et al. 2018) applies currently a compulsory 1 Hz high pass filter), and line noise removal (except ADJUST, Mognon et al. 2011). The Batch Electroencephalography Automated Processing Platform (BEAPP, Levin et al. 2018) and Automagic have included the PREP pipeline (Bigdely-Shamlo et al. 2015), which offers standardized early-stage EEG processing, includes sophisticated methods to remove line-noise (using Clean Line, Mullen 2012) and robust average referencing by iteratively detecting and interpolating artifactual channels. The Clean Line line-noise removal is also available in the automatic pre-processing pipeline for EEG analysis (APP) (da Cruz et al. 2018).

For artifact correction most pipelines have implemented an independent component analysis (ICA) (except Autoreject Jas et al. 2017). The majority of these pipelines (including Automagic) use MARA (Winkler, Haufe, and Tangermann 2011; Winkler et al. 2014) to automatically identify bad independent components. MARA is based on a supervised machine learning algorithm using six features from spatial, spectral and temporal domains. In addition to that, the BEAPP and HAPPE toolboxes allow to use wavelet-enhanced ICA. ADJUST uses a different set of spatial and temporal features to identify bad components. Only Automagic and partially Computational Testing for Automated Preprocessing (CTAP) (Cowley, Korpela, and Torniainen 2017; Cowley and Korpela 2018) (only EOG) offer additional artifact correction methods, such as EOG regression and robust PCA.

The CTAP toolbox also offers functions to optimize the different preprocessing methods by allowing the user to compare different pipelines. BEAPP is a platform for batch processing of EEG data that allows the creation of preprocessing pipelines with a variety of options that can be re-applied to new datasets, hence facilitating the exact methodological replication of studies.

Currently, next to Automagic only CTAP, and HAPPE offer quality metrics. The quality metrics of HAPPE are the number of epochs that survive epoch-wise artifact rejection after preprocessing, the number of independent components (IC) that are rejected as artifact components, and the retained variance of the EEG data after IC rejection. CTAP yields the percentage of bad channels, bad segments, bad epochs and bad components.

## 4 Validation

### 4.1 Methods

Automagic utilizes mainly preprocessing methods that have been previously published and most of the methods have been validated (Winkler, Haufe, and Tangermann 2011; Winkler et al. 2014; Bigdely-Shamlo et al. 2015; Mullen 2012; Parra et al. 2005). The main aim of this section is to validate the effect of *combining* methods on the resulting preprocessed data to provide recommendations on the choice of preprocessing steps when using Automagic.

The goal of preprocessing is to separate and clean artifacts (i.e. signal of no interest) from physiological signals of interest; hence, our validation aims to delineate to what extent noise can be separated from the signal of interest. Since it is not possible to determine the ground-truth for the signal of interest (i.e. EEG signal with neuronal source) because the neural sources cannot be fully determined by the EEG signal (i.e. the inverse problem), often validations are performed by simulating EEG with known sources of noise. However, it is not trivial to model the variability of artifacts between subjects that is typically observed in real data. Hence, we follow an indirect, yet more realistic validation strategy. We will later provide details but first outline the basic idea behind this strategy: Considering the EEG of one person, we assume that the underlying neuronal activity during rest is relatively similar during periods of time in which artifacts occur and periods of time with clean EEG. The EEG signals in the clean and artifact-contaminated time windows should, to a great extent, differ due to artifactual noise and not due to underlying differences in neuronal activity; hence, a valid preprocessing method should remove the artifactual noise and increase the similarity of key characteristics of periods of time where artifacts occur and periods of time without artifacts.

To test this, we used 52 EEG datasets (128 channels Netamps system (Electrical Geodesics, Eugene, Oregon), Cz recording reference, sampling rate: 500 Hz, 1 Hz high-pass filtered) of 26 healthy elderly subjects (mean age 71.5, SD ±5.1) during a passive resting task with opened eyes. The data and code that have been used are available (CC-By Attribution 4.0 International) online https://osf.io/5e74x/. The study was approved by the ethics committee of the canton of Zurich, Switzerland (BASEC-Nr: 2017-00226) and informed consent was obtained from each subject. The channels that were placed in the most ventral ring of the cap (E32, E48, E49, E56, E63, E68, E73, E81, E88, E94, E99, E107, E113, E119) were discarded. EOG channels were E1, E32, E8, E14. E17, E21, E25, E125, E126, E127, E128.

The data of each subject, consisting of five 16 s epochs (=40,000 time frames (TFs)), were first scanned for artifacts by identifying the TFs in which the standard deviation across channels is larger than 25 µV. Then, the EEG was epoched into 2 s segments. If a TF with a standard deviation greater than 25 µV was within an epoch, the epoch was marked as a bad epoch (i.e. epoch with artifact); otherwise marked as a good epoch (i.e. epoch without artifact).

Good and bad epochs were then concatenated to separate datasets. To avoid datasets with an exceedingly small number of time points, datasets with fewer than 10,000 TFs of either good or bad epochs were discarded for further analysis. To make the good and bad datasets of a subject comparable, the length of the dataset with more TFs was reduced to the length of the dataset with shorter length, by randomly choosing the corresponding number of epochs. This resulted in 52 datasets (for each subject one good and one bad dataset) with on average 22,000 TFs (min: 10,000 TFs, max: 30’000 TFs, std = 6,112 TFs).

In a next step, we set up and ran 18 different variations of preprocessing in Automagic. Although theoretically 32 potential different variations of preprocessing would be possible in Automagic, some combinations are not applicable in practice. For example, ICA (MARA) with bad channels interpolated would cause a processing error due to EEG rank deficiency and hence ICA alone cannot be used (e.g. (Ullsperger and Debener 2010). Thus, our validation included 18 reasonable combinations of preprocessing (see Figure 3, panel A) in Automagic. In a first set of variations, we preprocessed the datasets by only interpolating bad channels that were identified by PREP and/or clean_rawdata(), using the respective default settings (see section Bad channel identification and Bad channel interpolation). These bad channel interpolating methods were then combined with artifact correction methods of either MARA or rPCA and with a combination of EOG regression (with their default settings, see section Artifact correction). An additional validation analysis for event related potential (ERP) data with the same validation analysis idea can be found in Appendix B. The data and code for the ERP validation analysis can be found on OSF (https://osf.io/5e74x/).

**Figure 3.**
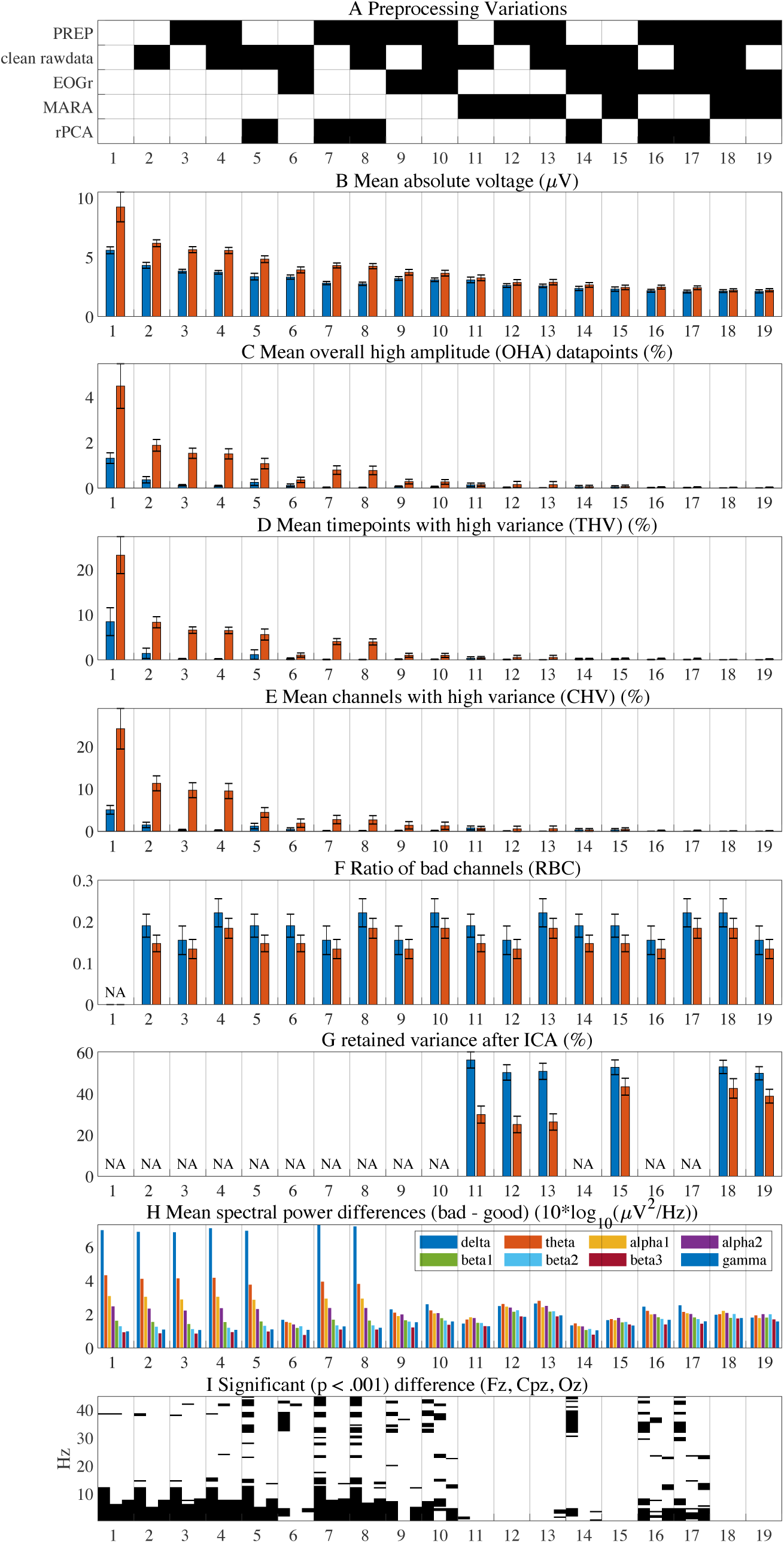
Validation results. Panel A shows the combinations of selected (black) preprocessing methods. The first column of the figure reflects the quality measures of the EEG data before the preprocessing. The combinations are sorted with respect to mean absolute voltage shown in panel B with blue bars for the good datasets and red bars for the bad datasets; hence, the order indicates how much signal is subtracted by the preprocessing variations. Panels C-E depict the results of applying the quality measures of overall high amplitude data points (OHA), time points with high variance (THV), and channels with high variance (CHV) (see section Quality measures for details on the calculation). Panel F shows the ratio of bad channels, as identified by the bad channel identification methods. In panel G the percentage of retained variance after the back-projection of non-artifactual ICs in MARA is shown. The difference between the spectra power in frequency bands of the bad minus the good datasets is shown in panel H. Panel I illustrates in which frequencies the power differs significantly (black bars: p < 0.001) in the channels Fz, Cpz, and Oz (each row reflects one channel). The error-bars reflect the standard errors.

### 4.2 Results

#### 4.2.1 Quantitative characteristics

The results of the preprocessing are summarized in Figure 3. The quality measures of the good and bad datasets were calculated and serve as comparison of the extent to which artifacts are present in the data before and after the respective preprocessing variation. Before preprocessing (Figure 3, leftmost column), the bad epochs show substantially more noise, (reflected in higher signal amplitudes) in all quality measures compared to the good epochs. In general, the more preprocessing methods are applied, the more noise and arguably overall signal of interest (see Figure 3 panel B) is removed. Thus, from this information alone it is not possible to assess an optimal preprocessing variation that balances removal of noise and signal. We will return to this later.

Considering the ratio of identified bad channels, clean_rawdata() detected on average 19% bad channels in the bad datasets and 15% in the good datasets, compared to fewer bad channels identified PREP (15% in bad datasets, 13% in good datasets). The combination of bad channel identification methods together yielded 22% of bad channels in the bad dataset and 18% of bad channels in the good datasets. Therefore, both bad channel identification pipelines identify to a great extent similar but not identical bad channels.

Panel G shows the percentage retained variance after MARA. This metric shows how much variance is retained after back-projecting the non-artifactual (clean) ICs. Note that retained variance is influenced by how clean the data are when entering the ICA of MARA. Therefore, more retained variance does not *per se* indicate less noisy data. If MARA is used without a preceding EOG regression, substantially more variance is removed from the data in the bad datasets (on average 52% retained variance in good datasets, 27% retained variance in bad datasets) whereas this difference is substantially smaller when MARA is used after EOG regression (on average 52% retained variance in good datasets, 42% retained variance in bad datasets). This indicates that the EOG regression and MARA may to some extent remove similar artifacts.

In addition to these rather global quality measures, we computed power spectral densities (PSDs) for the data before preprocessing (high-pass filtered at 1 Hz) and the 18 variations of preprocessing, for a frequency range of 1 Hz to 45 Hz (using the spectopo() function of EEGLAB with 2 times oversampling) for the channels Fz, Cpz, and Oz. Panel (H) shows to what extent the good and bad PSDs differ when aggregating spectral power bands (mean of “Fz”, “Cpz”, “Oz”, delta: 1.5 - 6 Hz, theta: 6.5 - 8 Hz, alpha_1_: 8.5 - 10 Hz, alpha_2_: 10.5 - 12 Hz, beta_1_: 12.5 - 18 Hz, beta_2_: 18.5 - 21 Hz, beta_3_: 21.5 - 30 Hz, gamma: 30 - 45 Hz). It becomes evident that the bad datasets have overall higher PSDs compared to the good datasets. Considering the different preprocessing variations, those with bad channel interpolation only and bad channel interpolation in combination with rPCA, leave large differences between good and bad data in lower frequency bands. The differences are substantially smaller when at least EOG regression or MARA is applied before.

To further quantify these findings, we statistically compared the “good” and “bad” PSDs of each subject. Recall that a valid preprocessing method should remove the artifactual noise and remove differences in PSDs in the good and bad epochs of subjects. To test this, we used paired-permutation tests: For each frequency bin separately, we calculate the difference in the frequency power between the “good” and “bad” dataset in each subject and sum these differences. This “real” sum of differences reflects the effect size of interest, which is compared to a distribution of 10’000 sums of difference values from pairwise randomly shuffled good and bad datasets. The ratio of instances in which the “real” sum of differences is smaller than the 10’000 randomly shuffled sum of differences, reflects the empirical probability p that the “real” difference would also have been detected by chance. Hence, a small p value in a certain frequency bin can be interpreted as that “good” and “bad” datasets differ with a small probability that this effect would have been observed by mere chance. The permutation tests are performed for each frequency bin (1 - 45 Hz) for the channels Oz, Cpz and Fz. The results of these tests are illustrated in Figure 3 panel I, where the frequency bins (x-axis), in which the PSD differs significantly (p < 0.001) between the good and bad datasets, are drawn in black for the channels Oz, Cpz and Fz (each channel in one row) and for each preprocessing variation. The results suggest that after applying any bad channel rejection method in combination with artifact correction using MARA, no significant differences in the PSD across the frequencies of 1 - 45 Hz are observable. In contrast, using bad channel rejection only or a combination thereof with rPCA and EOG regression leaves substantially more frequency bins that differ significantly.

#### 4.2.2 Qualitative characteristics

To assess to what extent signal (and not only artifacts) are removed, we consider qualitative characteristics of the PSDs. Figure 4A shows PSDs in the channels Fz, Pcz and Oz with only 1 Hz filtered data. Figure 4B shows the PSD after having applied PREP, Figure 4C PREP and EOGr, and Figure 4D PREP and MARA. The full set of plots of all variations are shown in Appendix A.

**Figure 4.**
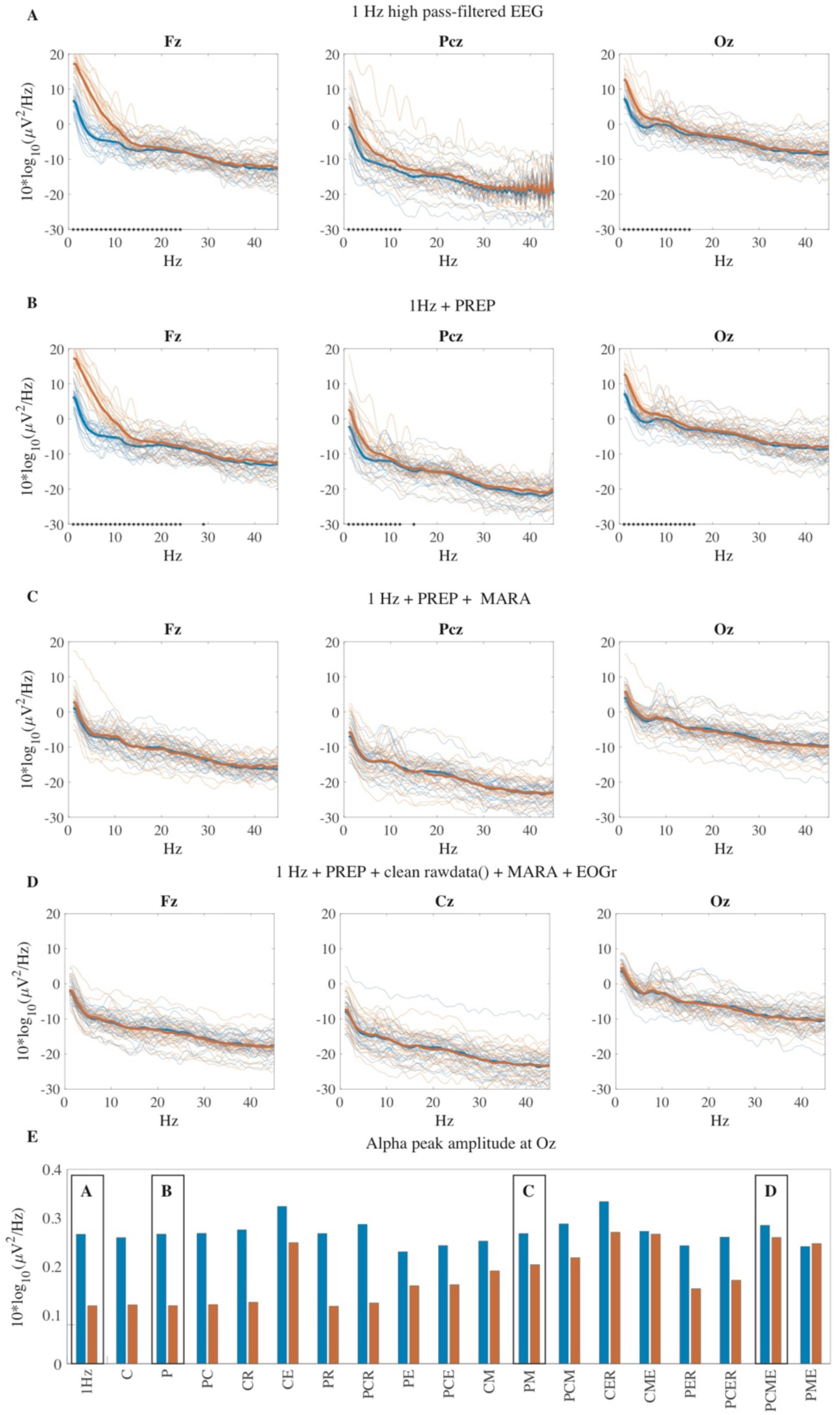
PSDs with different preprocessing variations. The PSDs of good (blue lines) and bad (red lines) datasets are shown for the channels Fz, Pcz, and Oz. The asterisks indicate significantly (p < 0.001) differing PSD at the specific frequency bin. Panel E shows the alpha peak power in Oz in good (blue) and bad (red) data after the preprocessing variations (in identical order as in Figure 2). The black boxes point to the respective preprocessing variation shown in panels A-F above. (P = PREP, C = clean rawdata(), M = MARA, R = rPCA, E = EOGr)

It is evident that only applying bad channel identification (i.e. PREP and clean_rawdata()) and interpolation leaves substantial differences between good and bad datasets in frequency ranges from 1 to 20 - 30 Hz (see also Appendix A. Figure 1 B-D). Considering the different combinations of artifact correction methods (i.e. EOGr, MARA, rPCA) and bad channel identification methods, our analysis demonstrates that MARA is able to correct for most artifacts in contrast to EOGr and rPCA, which leave substantial differences between good and bad datasets (see also Appendix A. Figure 1 E-S). Furthermore, the results suggest that a combination of PREP (or clear_rawdata()) and MARA is sufficient to remove (almost) all significant differences in the PSDs.

To estimate to what extent the preprocessing variations balance the removal of noise compared to retaining the signal of interest, we consider the characteristic alpha peak in Oz. We estimated the alpha peak power amplitude using the method by (Haller et al. 2018), where we calculated the mean alpha peak power from the mean (across subjects) PSD in the frequency range of 8.5 Hz to 12 Hz for each preprocessing variation. Even though the amplitude of the alpha peak is smaller with opened eyes compared to closed eyes (Klimesch 1999), the results show that before the preprocessing the alpha peak power is more than double as high in the good datasets compared to the alpha peak power in the bad datasets (see Figure 4E). With additional preprocessing, the alpha peak power in Oz in the bad datasets increases (Figure 4E, red bars) whereas the alpha peak in the good dataset (blue bars) does not substantially decrease, as it would if the signal of interest was removed by the preprocessing. This suggests that in the preprocessing (for instance using a variation of bad channel identification) MARA (and EOGr) removes to a great extent artifactual signal and not signal of interest (i.e. signal in the alpha band).

## 5 Discussion

The number of large-scale studies using EEG is limited compared to large-scale MRI and fMRI studies. As we argue, this is not because the EEG signals offers less informative insights into brain (dys-)functioning, but is manifested mainly in two unresolved issues. First, the quality of EEG recordings often substantially differs between subjects and currently there is no objective way of quantifying the quality and hence these differences. This renders the typically inevitable process of excluding subjects from analyses due to exceeding contamination of artifacts highly subjective. This not only impedes a precise methodological replication but also seduces researchers into trying out various combinations (i.e. P-hacking) and thereby further decreasing the chance of replication. The second reason is technical. Currently no software solution exists that controls and manages the preprocessing of ongoing and dynamically growing studies. To address these challenges, we developed Automagic, an open-source MATLAB toolbox that wraps currently available preprocessing methods and offers standardized quality assessment for growing studies. We introduced the functionality of Automagic and investigated the effect of applying combinations of methods on a sample of resting EEG data. This validation revealed that a combination of applying a pipeline of bad channel identification algorithms and MARA, an ICA-based artifact correction method, is sufficient to reduce a large number of artifacts.

There are several limitations to the presented validation and also to the use of Automagic. First, the ratio between the duration of the data and the number of channels used for the validation is rather low and may be suboptimal for an ICA decomposition. However, we believe that the result of the validation—namely, that a combination of MARA and a bad channel identification pipeline successfully removes noise while preserving the characteristics of resting EEG PSDs (e.g. alpha peak)—still holds. With more data, the ICs would be estimated more robustly and hence the results should improve. Yet, it is also possible that the performance of correcting artifacts with the rPCA and EOGr improves with more data.

The results of the validation are furthermore limited to testing the effects of preprocessing on the PSDs of resting EEG with eyes open. Other signals of interests (such as characteristics of event related potentials (ERP), features in time-frequency decomposition, source localization and connectivity measures) may favor different combinations of preprocessing and should be tested on the respective datasets. We chose resting EEG with open eyes because we aimed to use realistic EEG data with a similar number of clean segments and segments of EEG that were clearly contaminated with artifacts. Under the assumption that the true underlying neuronal activity is similar in the artifactual and clean epochs, we assessed the effectiveness of the methods by considering the extent of increase in the similarity of key characteristics of the good and bad datasets.

## 6 Conclusion

We introduced Automagic an open-source MATLAB toolbox that incorporates currently available EEG preprocessing methods and offers new objective standardized quality assessment for growing studies. Automagic fosters transparency and data sharing in EEG by complying with the BIDS EEG standard and providing detailed logging of all performed processes. With this we hope to facilitate future large scale EEG studies to advance the replicability of EEG research and the detection of EEG-based biomarkers.

## Author contributions

A.P, A.B. and N.L. developed the software. A.P. and N.L. designed the validation. A.P. and N.L. analyzed the data. A.P. and N.L. wrote the paper.

## Acknowledgements

We thank Dominik Güntensperger, Sabine Dziemian, Marius Tröndle, Martyna Plomecka & Christian Pfeiffer for extensively testing Automagic and for assistance with manuscript revision.

## Appendix A

**Figure 1.**
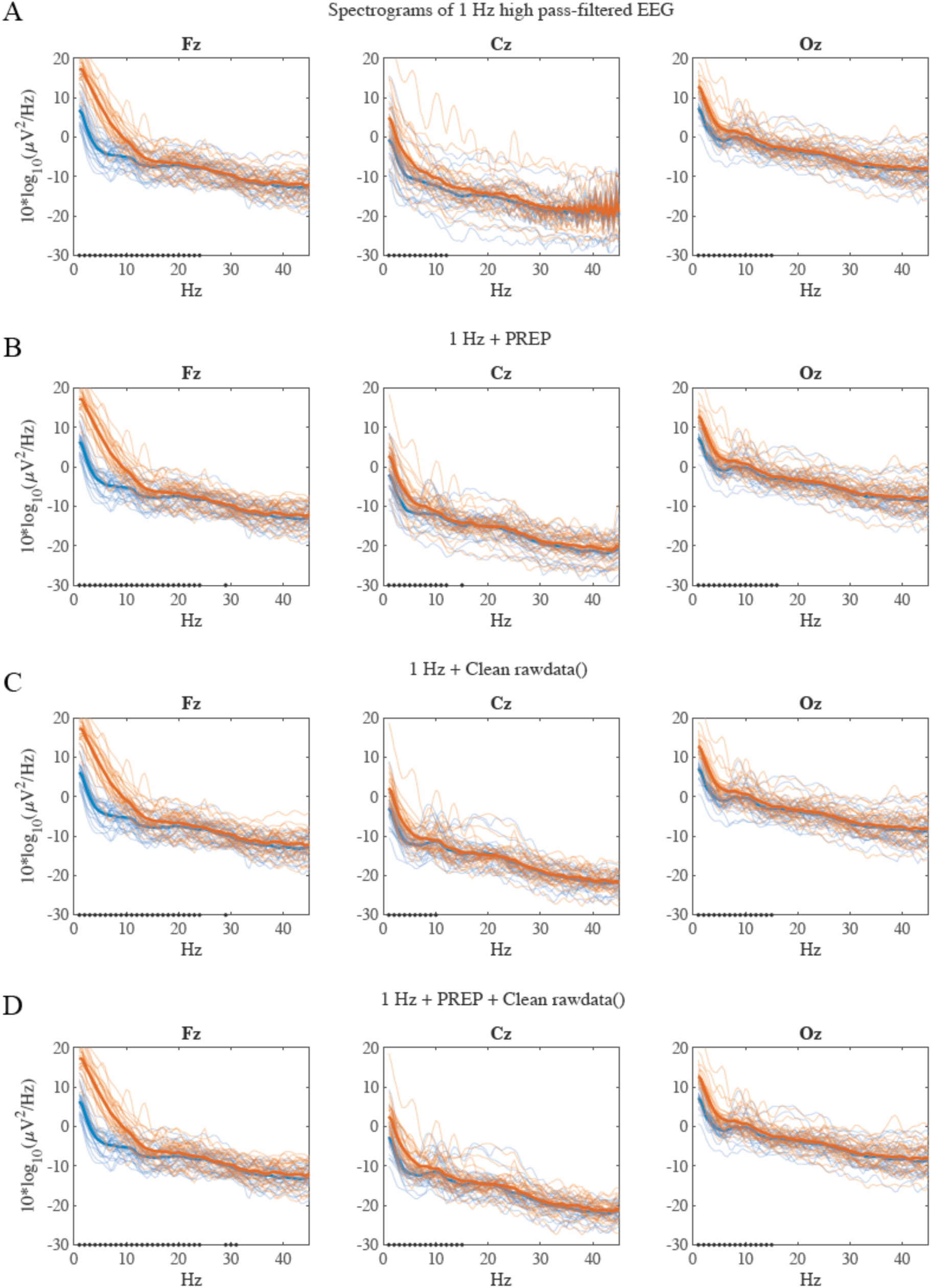
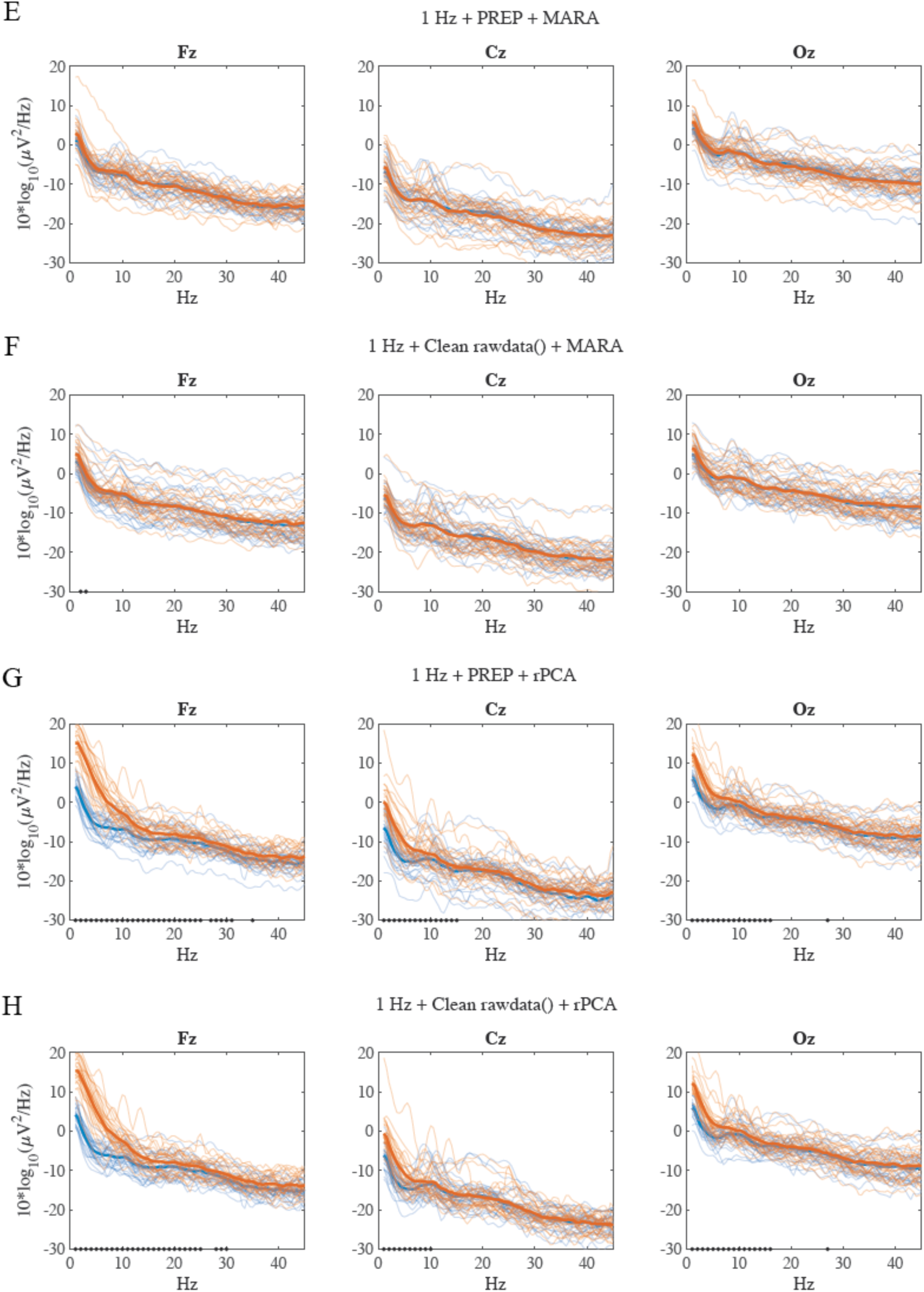
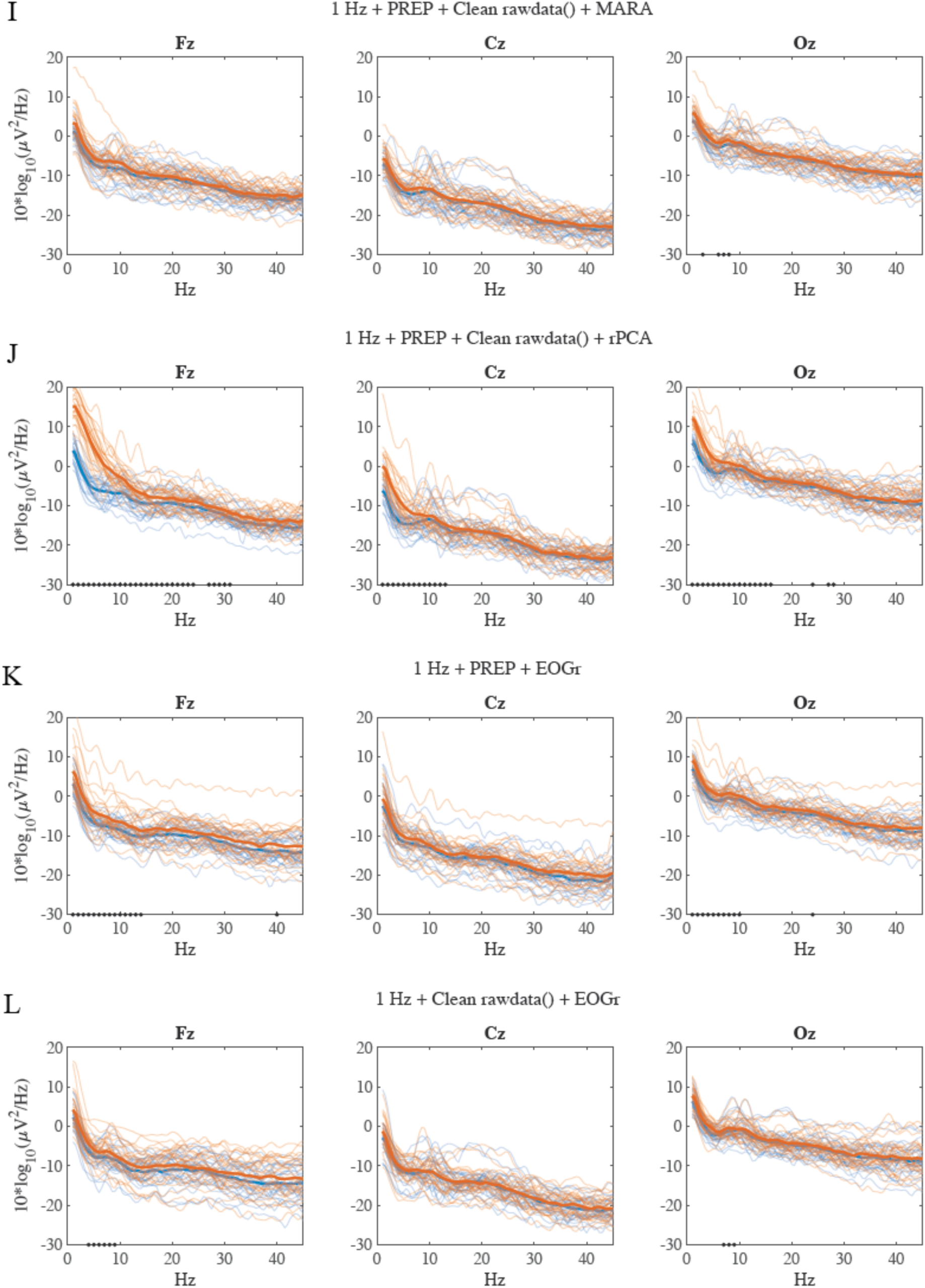
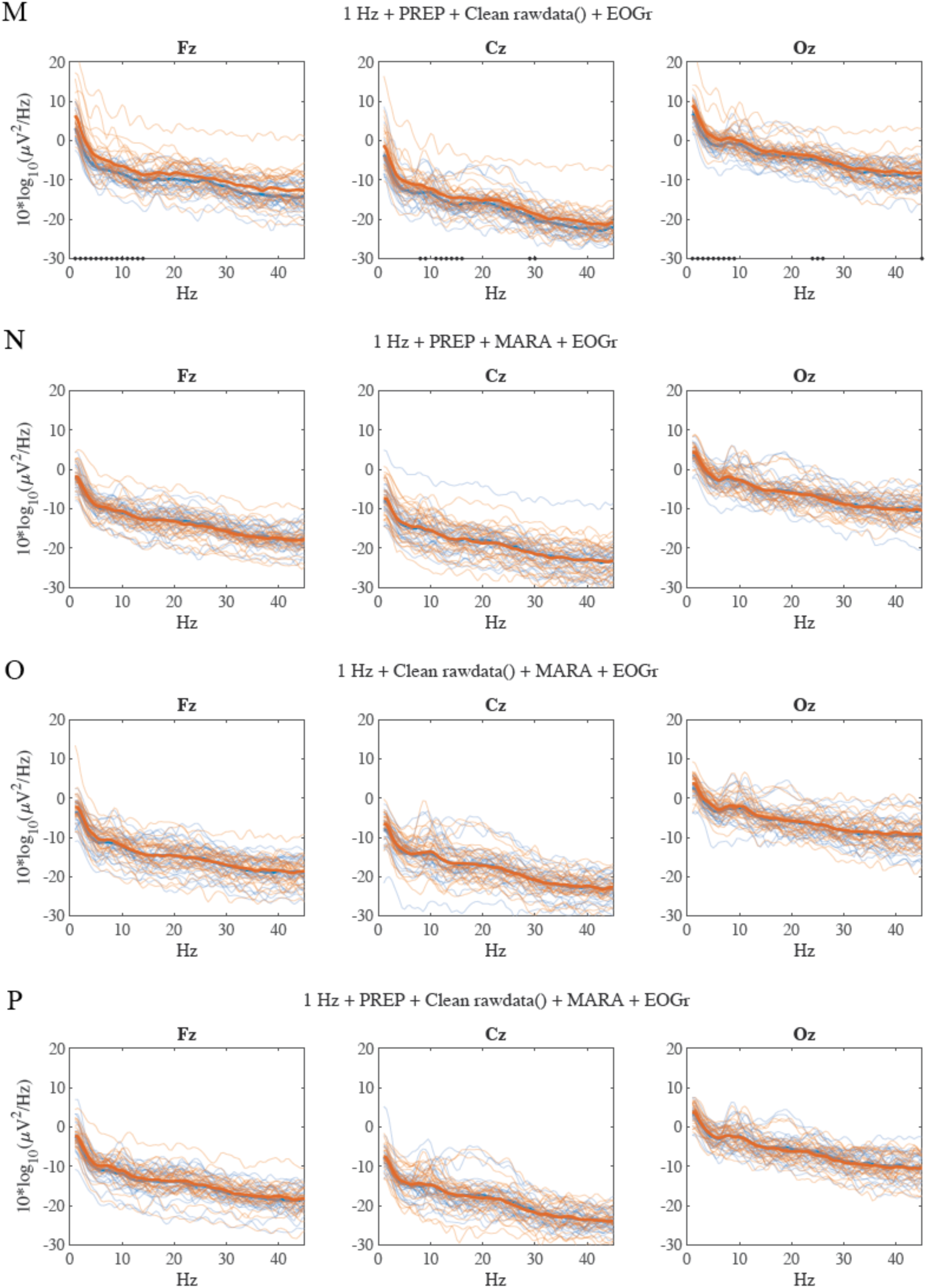
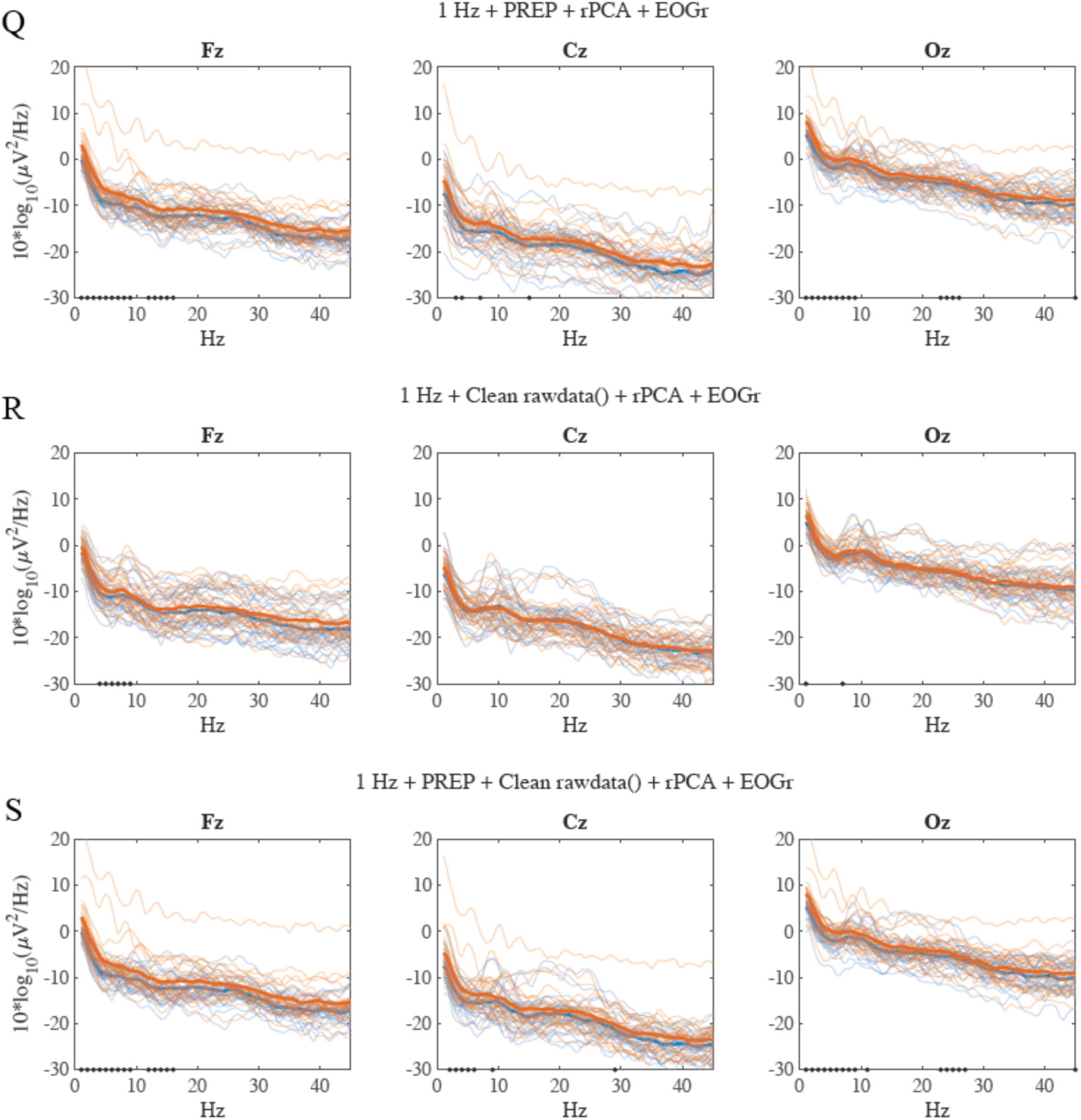

## Appendix B

### Event Related Potentials (ERP) Validation Analysis

The data and code for the Event Related Potentials (ERP) validation analysis can be found on OSF (https://osf.io/5e74x/).

### Methods

To test whether Automagic is also valid for Event Related Potentials (ERP) data we used a similar validation approach as for the resting state EEG data. Signals in the clean and artifact-contaminated time windows should, to a great extent, differ due to artifactual noise and not due to underlying differences in neuronal activity; hence, a valid preprocessing method should remove the artifactual noise and increase the similarity of key characteristics of an artifactual ERP and artifact-free ERP.

The ERP validation analysis was performed on a publicly available dataset of (Wakeman and Henson 2015). This dataset was collected within the OpenNeuro project (https://www.openneuro.org). accession #:ds000117. 16 subjects with an age range 23–37 years successfully completed the experiment. Each subject performed six blocks of a passive perceptual task on pictures of familiar, unfamiliar and scrambled faces. For the current validation analysis, we only analyzed the trials with unfamiliar faces with the goal to extract the classic face-sensitive N170 ERP component. A total of 296 trials was available for each subject. Each trial started with a fixation cross for a random duration between 400 and 600 ms, after which the critical unfamiliar face was presented for a duration between 800 and 1000 ms and an intertrial interval of 1700ms. Participants were instructed to fixate centrally throughout the experiment. For further details about the stimuli and experimental design please see (Wakeman and Henson 2015). EEG was recorded in a light magnetically shielded room using a Elektra Neuromag Vectorview 306 system (Helsinki, FI) and a 70 channel Easycap EEG cap (based on EC80 system) system (Wakeman and Henson 2015). Data were acquired at an 1100 Hz sampling rate with a low-pass filter at 350 Hz and no high-pass filter. The EEG reference electrode was placed on the nose, and the common ground electrode was placed at the left collar bone. Two bipolar electrodes were used to measure vertical and horizontal electrooculography (VEOG and HEOG).

For our validation analysis, data were resampled to 250 Hz and high-pass filtered (0.5 Hz). Subsequently, the data were segmented (−100ms to 800ms) and baseline-corrected. Similar to the resting EEG validation analysis, we have first scanned each epochs for artifacts by identifying the time frames (TFs) in which the standard deviation across channels is larger than 25 µV. If a TF with a standard deviation greater than 25 µV was within an epoch, the epoch was marked as a bad epoch (i.e. epoch with artifact); otherwise marked as a good epoch (i.e. epoch without artifact). To avoid subjects with an exceedingly small number of epochs, subjects with fewer than 30 of either good or bad epochs were discarded for further analysis. To make the good and bad datasets of a subject comparable, the number of epochs was reduced to the number of epochs of the dataset with smaller number of epochs, by randomly choosing the corresponding number of epochs. However, the index of the selected epochs was stored to compare the same good and bad trials across the different preprocessing pipelines. This resulted in 10 subjects with at least 30 good and bad trials (an average of 78 trials per subject). In a next step, we set up and ran 18 different variations of preprocessing in Automagic. These are the identical 18 combination of preprocessings as for the resting EEG validation analysis. The input for each of the 18 Automagic variation was the continuous data, including all trials (the selected bad and good trials and the not selected trials). After preprocessing the data with each of the 18 combinations, we have segmented and baseline-corrected the data for each subject and preprocessing variation again as described above. Finally, we have computed ERPs for the initially defined good and bad epochs, resulting in a “good” and “bad” ERP for each subject and preprocessing pipeline. Identical to (Wakeman and Henson 2015), the ERP was obtained from a right parieto-occipital electrode (‘EEG065’). Wakeman and Henson (2015) have demonstrated in the same data set a negative deflection peaking around 170 ms (‘N170 component’). Several studies have demonstrated that human faces elicit a evident N170 component (Allison et al. 1999; Bentin et al. 1996). We have additionally analyzed an earlier positive deflection around 100 ms (‘P100 component’), which is the major component of the visual evoked potential (VEP) and very reliable between individuals (Emmerson-Hanover et al. 1994; O’Shea, Roeber, and Bach, 2010). Subsequently we compared the good and bad epochs and consequential good and bad ERPs across the different Automagic preprocessing variations.

### Results

The results of the preprocessing are summarized in Figure 1 of Appendix B. The quality measures of the good and bad datasets were calculated and serve as comparison of the extent to which artifacts are present in the data before and after the respective preprocessing variation. Before preprocessing (Appendix B Figure 1, leftmost column), the bad epochs show substantially more noise, (reflected in higher signal absolute amplitudes across the ERP) in all quality measures compared to the good epochs. In general, the more preprocessing methods are applied, the more noise and arguably overall signal of interest (see Figure 1 of Appendix B panel B-E) is removed. However, from this information alone it is not possible to assess an optimal preprocessing variation that balances removal of noise and signal (see discussion of resting EEG data validation). Considering the ratio of identified bad epochs (see panel F), the only high-pass filtered data (0.5 Hz) identified 24% of bad trials (across all 296 trials). While bad channel interpolation methods reduced the ratio of bad epochs to an average of 9%, additional PCA and ICA artifact correction reduced the ratio of bad trials to an average of 1%. In addition to these rather global quality measures, we computed the signal-to-noise ratio (SNR) for N170 and P100. The SNR was computed as the ratio of the peak amplitude for N170 or P100 and the averaged pre-stimulus noise (De Vos, Gandras, and Debener 2014; Oliveira et al. 2016). Pre-stimulus noise was defined as the root-mean square of the period from -100 to 0 ms and was calculated for each single trial (De Vos, Gandras, and Debener 2014; Oliveira et al. 2016). The SNR for N170 and P100 were computed for the data before preprocessing (high-pass filtered at 0.5 Hz) and the 18 variations of preprocessing. Panel (G) for the SNR N170 component and panel (H) for the SNR P100 component shows to what extent the good and bad epochs differ. It becomes evident that the bad ERP’s have overall lower SNR compared to the good ERP’s. Considering the different preprocessing variations, those with bad channel interpolation only and bad channel interpolation in combination with rPCA, display a similar SNR for the N170 and P100 component as the SNR for the data before preprocessing. However, when ICA MARA is applied, the SNR increases for up to 30%. Recall that a valid preprocessing method should remove the artifactual noise and remove differences in amplitude in the good and bad ERPs, resulting in a high SNR, which is similar between good and bad ERPs.

**Figure 1.**
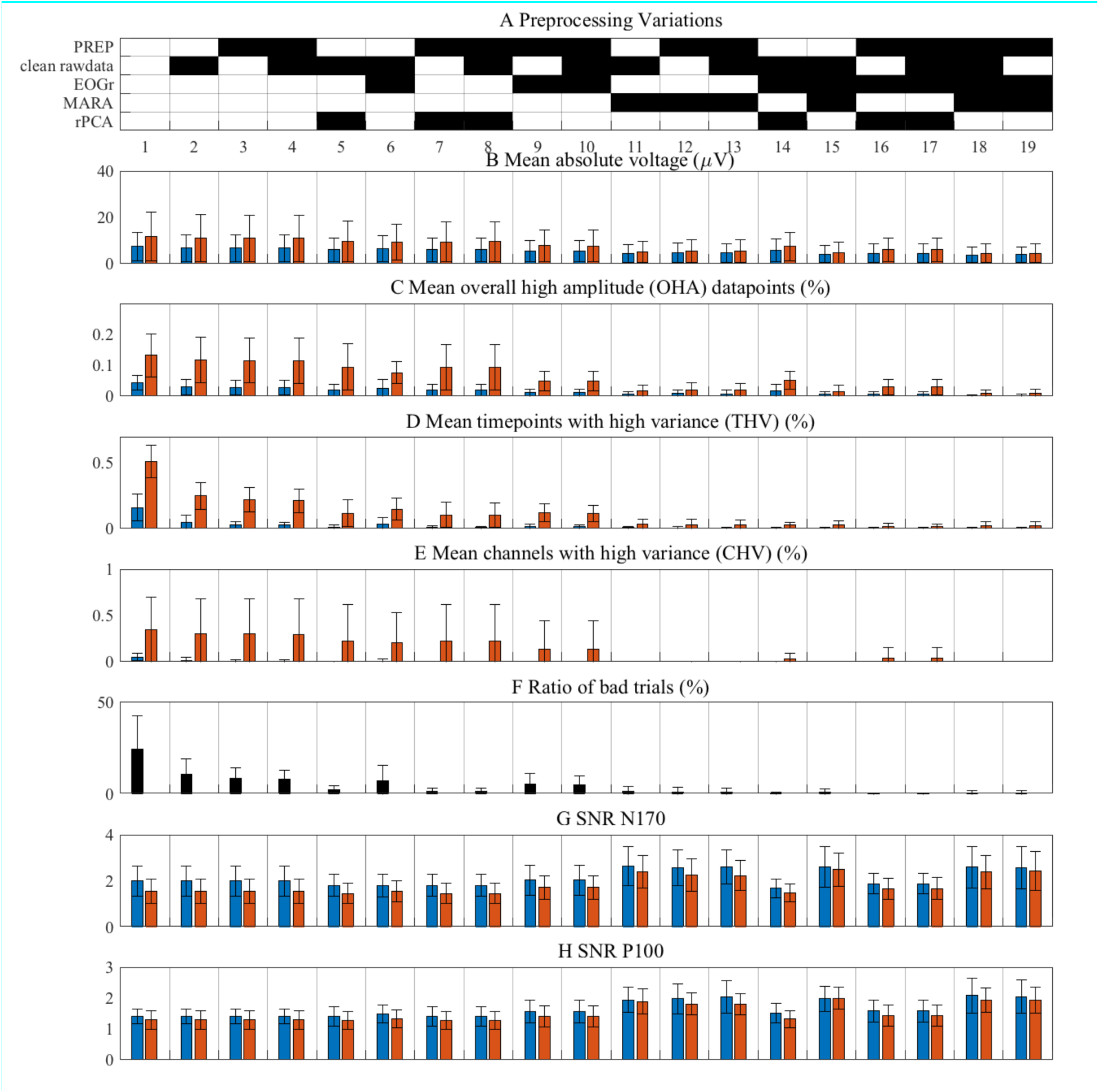
ERP Validation results. Panel A shows the combinations of selected (black) preprocessing methods. The first column of the figure reflects the quality measures of the EEG data before the preprocessing. The blue bars indicate the good datasets and red bars represent the bad datasets; hence. Panels C-E depict the results of applying the quality measures of overall high amplitude data points (OHA), time points with high variance (THV), and channels with high variance (CHV) (see section Quality measures for details on the calculation). Panel F shows the ratio of bad trials, as identified by a time frame with a standard deviation greater than 25 µV was within an epoch. In panel G the signal-to-noise ratio (SNR) for the N170 component is depicted. Panel H exhibits the SNR for the P100 component. The error-bars reflect the standard errors.

To further quantify the differences between good and bad ERP, we statistically compared the “good” and “bad” ERP time point by time point. We used the identical paired-permutation tests as in the resting EEG validation analysis, but replaced the frequency bins with the time points. The permutation tests are performed for each timepoint for the channel ‘EEG065’. The results from the paired-permutation tests are illustrated in Figure 2 of Appendix B. The significant time points are indicated as asterisks along the x-axis (time). Moreover, Figure 2 displays the bad and good ERP’s for each preprocessing variation. The results suggest that after applying any bad channel rejection method in combination with artifact correction using MARA, no significant differences in the amplitude for the time segment between 170-200ms are observable anymore. In contrast, using bad channel rejection only or a combination thereof with rPCA and EOG regression leaves substantially more time points between 180-200ms that differ significantly. Furthermore, late time segments (>400ms) show less significant differences between the good and bad ERP’s when ICA MARA in combination with a bad channel detection and EOG regression has been conducted. In summary, the ERP validation revealed that a combination of applying a pipeline of bad channel identification algorithms and MARA, an ICA-based artifact correction method, is sufficient to reduce a large number of artifacts.

**Figure 2.**
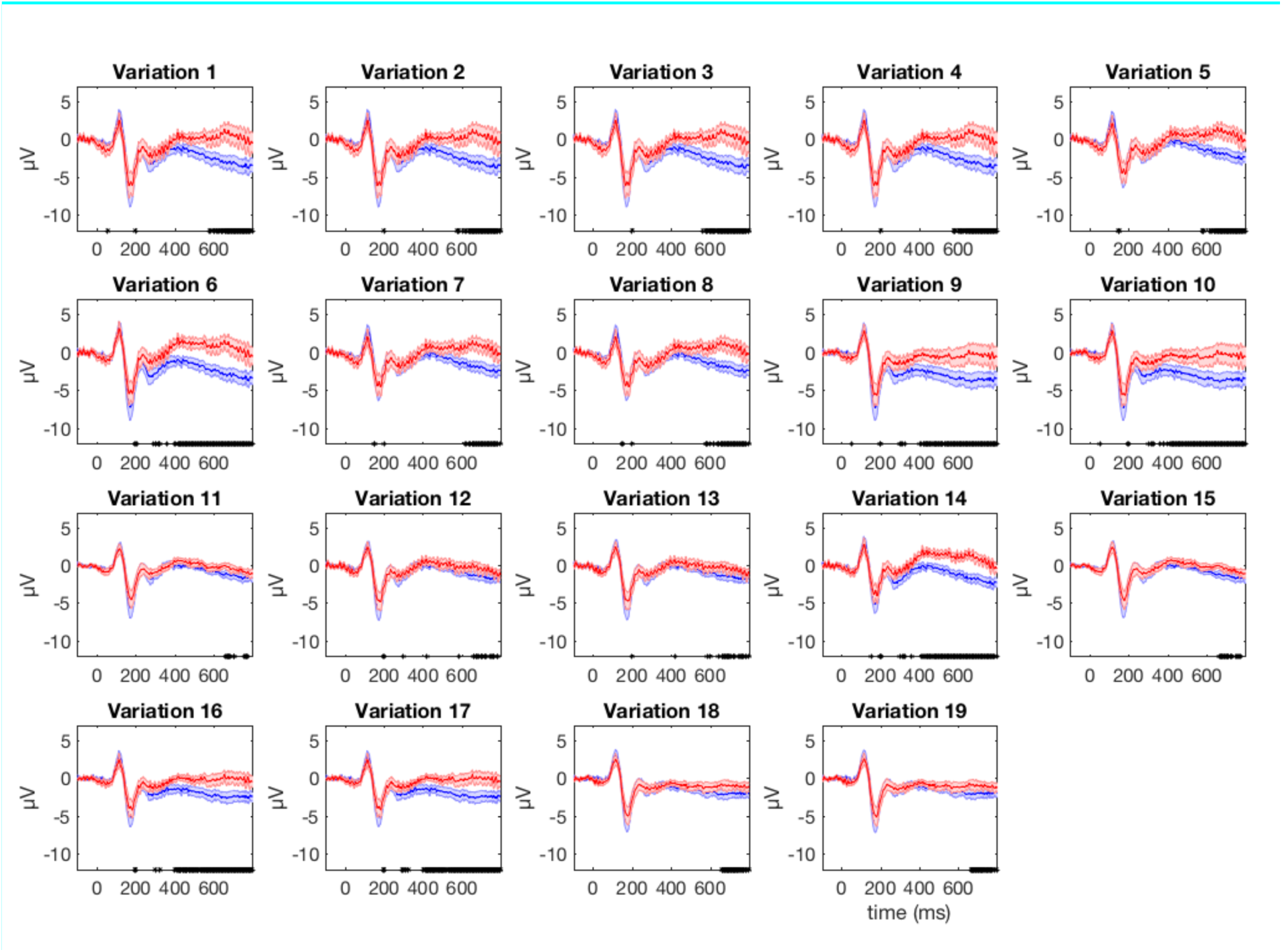
ERPs with different preprocessing variations. The ERP’s of good (blue lines) and bad (red lines) datasets are shown for the right parieto-occipital electrode (EEG065). The standard error is shown as shaded area around the mean. The asterisks along the x-axis indicate significantly (p < 0.005) differing amplitude at the specific time frame as a result from the paired-permutation statistics. The numbering of the variations is identical to the numbering of the variations in Figure 1 of the Appendix B.

## Footnotes

1 The visual classification of EEG has been shown to be moderately reliable (kappa = 0.75) and can be improved with training (kappa = 0.91) (Hatz et al. 2015).

2 Note that it is possible to skip bad channel identification, but it is advisable to identify and later exclude bad channels before applying artifact correction routines because methods such as the ICA applied in MARA or the rPCA may not work properly if bad channels are not excluded beforehand.

3 Note that changing the default settings of clean_rawdata() needs to be done in the respective functions.

4 Note that the data, which is the input to preprocessing, are not modified at any time.

5 EEGLAB functions can be directly applied to the loaded EEG structure. Alternatively, by starting EEGLAB and calling the command eeglab-redraw the EEG structure in MATLAB’s workspace becomes accessible in the EEGLAB GUI.

6 Users may have observed a reduced_*.mat file in the results folders. This file is a downsampled version of the EEG data and is used for faster displaying in the data viewer.

7 Note that this recommendation is based on data recorded with an 128 channels Netamps system (Electrical Geodesics, Eugene, Oregon) and may not apply to other recording systems.

